# Characterizing the polarization continuum of macrophage subtypes M1, M2a and M2c

**DOI:** 10.1101/2022.06.13.495868

**Authors:** TCL Oates, PL Moura, SJ Cross, K Roberts, HE Baum, KL Haydn-Smith, MC Wilson, KJ Heesom, CE Severn, AM Toye

## Abstract

Macrophages are vital components of the inflammatory response and exhibit phenotypical plasticity through active conversion between pro- and anti-inflammatory cell subtypes, a feature which can be reproduced in *ex vivo* culture. We employed a multifaceted approach utilizing proteomics, flow cytometry, activity assays and livecell microscopy imaging to characterize four cultured macrophage subtypes: unstimulated MØ, classically activated M1, alternatively activated M2a, and deactivated M2c macrophages. Whole cell proteomics identified a total of 5435 proteins, with >50% of these proteins exhibiting significant alterations in abundance between the different subtypes. This confirms that four distinct macrophage subtypes are induced from the same originating donor material through stimulation with specific cytokines. Additional surfaceome analysis revealed that M2c macrophages significantly upregulate pro-inflammatory markers compared to the MØ baseline and thus appear to be activated or primed to activate, similar to M1. Surface protein expression provided further subtype characterization, in particular distinguishing between the M2a and M2c macrophages.

We next explored the re-polarization capabilities of macrophages using dexamethasone, an anti-inflammatory glucocorticoid known to induce macrophage polarization towards the M2c de-activated phenotype. We show that activated M1 macrophages treated with dexamethasone for 48-hours upregulate the levels of CD163 and CD206, markers synonymous with a phenotypical shift from M1 to M2c yet retain key surface markers and display the functional phenotype of M1 macrophages. The observed repolarization of M1 pro-inflammatory macrophages provides a potential mechanism through which dexamethasone treatment improves COVID-19 prognosis and constitutes evidence of partial repolarization along the macrophage continuum. These proteomic and functional *ex vivo* macrophage datasets provide a valuable resource for studying macrophage polarity and the impact of dexamethasone on macrophage phenotype and function.

## 1 Introduction

Macrophages are specialized immune cells with broad functional heterogeneity. In addition to the detection and destruction of bacteria and viruses, macrophages produce a cocktail of cytokines stimulating a variety of immuno-pathologies, including Coronavirus Disease-19 (COVID-19)(Huang, Wang et al. 2020, Mehta, McAuley et al. 2020). An important feature of macrophages is their ability to exhibit plasticity in their phenotype, differentiating into pro- and anti-inflammatory subtypes in response to cytokine stimulation and local tissue environment (Roszer 2015). The first archetypal antimicrobial macrophage identified was induced by exposure to interferon-g (IFNg) secreted by lymphocytes; displaying oxidative metabolism and antimicrobial activity and were named ‘classically activated’ macrophages (Nathan, Murray et al. 1983). T helper cells, Th1 and Th2, exhibit distinct cytokine secretion profiles leading to the generation of macrophages with unique phenotypes. Th1 cells induce the pro-inflammatory ‘classically activated’ macrophages referred to as M1, whilst Th2 induce M2 macrophages that are ‘alternatively activated’ (Mosmann and Coffman 1989, Mills, Kincaid et al. 2000). Mantovani *et al*. further subdivided the M2 macrophages into M2a, M2b, M2c and M2d subtypes based on applied stimuli and the transcriptional changes induced (Mantovani, Sica et al. 2004). M2a macrophages are ‘alternatively activated’ induced by Interleukin-4 (IL-4), involved in clearance of apoptotic cells, modulating the pro-inflammatory response and wound healing (Stein, Keshav et al. 1992), whilst the M2c phenotype induced by glucocorticoids are considered anti-inflammatory ‘deactivated’ macrophages with a role in tissue remodeling, and are central to the erythroblastic island (Bessis 1958, Chasis and Mohandas 2008, Roszer 2015, Heideveld, Hampton-O’Neil et al. 2018). Macrophage subtypes are generally identified based on cell surface markers. M1 macrophages being defined by high expression of CD14, CD80, CD86 and CD16; M2a macrophages typically have lower expression of CD14 and CD16 and higher expression of CD206, and M2c macrophages comparatively express high levels of CD14, CD169, CD163 and CD206 (Mantovani, Sica et al. 2004, Gundra, Girgis et al. 2014, Raggi, Pelassa et al. 2017, Heideveld, Hampton-O’Neil et al. 2018). Importantly, beyond the dichotomy of classically and alternatively activated macrophages, further heterogeneity exists in macrophage populations, along with their ability to repolarize to adapt to tissue environments.

Our current understanding of macrophage polarity has been expanded beyond cell surface marker identification through proteomic investigations. The most comprehensive of these studies conducted by Court *et al*., focused on human *ex vivo* CD14^+^ derived macrophage polarity in environmental oxygen tension (Court, Petre et al. 2017). Heideveld *et al*. used TMT-labelled whole-cell proteomic comparison between dexamethasone stimulated and unstimulated human *ex vivo* CD14^+^ derived macrophages, highlighting a shift towards an anti-inflammatory subtype (Heideveld, Hampton-O’Neil et al. 2018). Finally, Becker *et al*. compared mouse M1 and M2 macrophages with dendritic cells, concluding that dendritic cells, classically activated, and alternatively activated macrophages were distinct entities and can be described as such from proteomic signatures (Becker, Liu et al. 2012). “Surfaceome” studies on fluorescence-activated cell sorted (FACS) classical and intermediate monocytes have also been conducted (Ravenhill, Soday et al. 2020). Importantly though, no study has compared proteomes and behavior of multiple human macrophage subtypes generated *ex vivo* from the same originating donor monocyte population to better understand the changes induced by macrophage plasticity.

COVID-19 is a respiratory disease causing a spectrum of symptoms from mild illness to death in addition to asymptomatic carriers. Functional and morphological differences have been noted in macrophages isolated from hospitalized COVID-19 patients (Huang, Wang et al. 2020, Mehta, McAuley et al. 2020). In addition to changes in size and granularity of the cells, a shift in macrophage polarization away from M1 macrophages via a decrease in CD14^+^/CD16^-^ expression has been noted within COVID-19 patient peripheral blood monocyte populations, whilst also observing an increase in CD80 and CD206 expression, suggesting that macrophage activation status plays a role within the disease (Zhang, Guo et al. 2020). Zhou *et al*. reported “interferon signaling ranked first for upregulated pathways in COVID-19” suggesting a role for M1 macrophages in particular (Zhou 2020). Dexamethasone, a glucocorticoid with anti-inflammatory and immunosuppressant effects, is proven to be successful in the treatment of COVID-19 patients requiring additional support from oxygen or mechanical ventilation reducing mortality by 29.3% and 23.3% respectively (Group, Horby et al. 2020). We, and others, have previously shown that activation of the glucocorticoid receptor induces differentiation of macrophages towards an M2c phenotype (Ehrchen, Steinmuller et al. 2007, van de Garde, Martinez et al. 2014, Falchi, Varricchio et al. 2015, Heideveld, Hampton-O’Neil et al. 2018, Hampton-O’Neil, Severn et al. 2020).

In this current work we comprehensively characterize four human *ex vivo* generated macrophage subtypes and their polarity, with a focus on semi-quantitative proteomics and functionality. Importantly, we demonstrate distinct expression patterns of proteins between the subtypes which is dependent on the macrophage phenotype and function. These include identifying significant differences between M2a and M2c macrophages where the literature does not frequently distinguish between the subtypes. We also explore the effects of dexamethasone exposure on the inflammatory M1 subtype, characterizing the effect of this steroid on macrophage polarization. Through this detailed dissection of macrophage biology, we improve understanding of macrophage polarity, providing novel datasets relevant to both COVID-19 and the macrophage research field.

## 2 Methods

### 2.1 Antibodies

Primary antibodies used: CD14 VioBlue (BD Pharminogen, Cat: 558121), CD16 FITC (Miltenyi Biotec, Cat: 130-113-392), CD80 FITC (BioLegend, Cat: 305205), CD86 PE (BioLegend, Cat: 305405), CD119 FITC (Miltenyi Biotec, Cat: 130-099-929), CD120a APC (BioLegend, Cat: 369905), CD121a (BD Biosciences, Cat: 551388), CD163 PE (Miltenyi Biotec, Cat: 130-097628), CD169 APC (Miltenyi Biotec, Cat: 130-098-643), CD192 PE (Miltenyi Biotec, Cat: 130-118-477), CD206 APC (BD Pharminogen, Cat: 550889), CD209 (BioRad, Cat: MCA2318T), CD210 PE (Miltenyi Biotec, Cat: 130-121-426) and ACE2 (R&D Systems, Cat: AF-933-SP). Secondary antibodies used: F(ab’)2-Donkey anti-Goat IgG cross absorbed PE secondary antibody (Thermo Fisher Scientific Cat: 31860), anti-mouse APC secondary antibody (Biolegend, Cat: 406610), anti-mouse Brilliant Violet 510™ secondary antibody (BioLegend, Cat:406621). Antibodies were used as per manufacturer’s instructions.

### 2.2 Peripheral blood mononuclear cell and CD14^+^ isolation

Peripheral blood mononuclear cells (PBMNC) were isolated from platelet apheresis blood waste (NHSBT, Bristol, UK) from anonymous healthy donors and those who had donated convalescent plasma with informed consent. Ethics approval for all experimental protocols was granted by Research Ethics Committee (REC 12/SW/0199 and 20YH/10168), and all methods were carried out in accordance with approved guidelines. PBMNC separation was performed using PBMC Spin Medium^®^ (pluriSelect Life Science) as described previously (van den Akker, Satchwell et al. 2010, Bell, Satchwell et al. 2013). Briefly, blood from apheresis cones was diluted 1:1 with Hanks balanced salt solution (HBSS, Sigma) containing 0.6% acid citrate dextrose (ACD) and layered over PBMC Spin Medium^®^. Samples were centrifuged at 400g for 30 minutes without brakes to generate a density gradient. CD14^+^ cells were isolated from PBMNCs using a magnetic micro-bead CD14^+^ kit (Miltenyi Biotec), LS columns (Miltenyi Biotec) and MidiMACSä separators (Miltenyi Biotec) as per the manufacturer’s instructions.(Heideveld, Hampton-O’Neil et al. 2018) CD14^+^ cells were stored frozen in 50% foetal calf serum (FCS, Gibco), 40% PBS with 10% DMSO (Sigma) in liquid nitrogen until required.

### 2.3 Macrophage ex vivo culture

Thawed cells were washed in PBS and resuspended at a density of 0.17×10^6^/mL – 0.33×10^6^/mL in RPMI 1640 (Gibco) supplemented with 10% FBS (Gibco), 25ng/mL macrophage–colony stimulating factor (wt/vol, M-CSF, Miltenyi Biotec), and penicillin/streptomycin at 100U/0.1mg per mL of media respectively (wt/vol; Sigma); with or without the inclusion of interleukin-4 at 20ng/mL (wt/vol, IL-4, BioLegend), interferon-g at 2.5ng/mL (wt/vol, IFNg, BioLegend) and dexamethasone 1mM/mL (wt/vol, Sigma), for differentiating macrophages. Cells were incubated at 37°C with 5% CO2. Full media changes were performed twice throughout the 7-day culture where any cells in suspension were collected via centrifugation at 300g and replaced in the culture along with the fresh media. Cells were harvested from the adherent macrophage culture via scraping and using a detaching buffer (10mM EDTA, 15mM Lidocaine in PBS) where required for flow cytometry analysis.

### 2.4 Flow cytometry

Analysis of cell surface markers by flow cytometry was performed using 0.5-1×10^5^ cells labelled either with directly conjugated antibodies or with a primary and secondary antibody combination. Antibodies were incubated for 30 minutes at 4°C. Data was collected using a MACSQuant flow cytometer (Miltenyi Biotec) and analyzed using FlowJo Version 10.7.

### 2.5 IncuCyte^®^ image acquisition and analysis

For imaging assays cells were seeded at 0.75×10^5^/well of a CELLSTAR^®^ 12 Well Plate on day 6 of culture and left to adhere overnight. The following day media was changed to remove suspension cells and images were captured using the IncuCyte^®^ SX1 Live-Cell Analysis System (Essen BioScience) using a 20x objective (0.45 NA) to capture 25 imaging fields for each donor every 20 minutes over a 12-hour time course. Videos were composed using the IncuCyte^®^ software and analyzed using a custom macro on Fiji (ImageJ). For MØ, M2a, and M2c subtypes individual cells were identified as ‘cells’ and tracked over time. M1 macrophages were observed to cluster during cultures, in which case the clusters were tracked as a single object due to the difficulty of tracking the single cells comprising the cluster. Both ‘cells’ and ‘clusters’ underwent the same analysis stream within the custom Fiji macro. Representative images from this analysis pipeline can be found in Supplemental Figure 1.

### 2.6 Imaging analysis pipeline

Cells were tracked from phase contrast videos using the MIA modular workflow plugin for Fiji.(Schindelin, Arganda-Carreras et al. 2012, Rueden, Schindelin et al. 2017, Cross 2019) First, videos were loaded using a JavaCV-based plugin for Fiji (Bradski 2000, Audet 2019, Cross 2020) with drift in the input videos corrected using a translation-only affine transform calculated using Fiji’s SIFT feature extraction plugin (Lowe 2004, Saalfeld 2009). For the purpose of cell detection, a copy of the raw video was made and converted to 16-bit without intensity scaling, then passed through a 2D local variance filter (σ = 5px) to enhance cells against the relatively homogeneous background. This video was binarized using the Otsu threshold method (Otsu 1979) and holes in the resulting binary image filled. The raw video was also sequentially passed through a difference of Gaussian filter (σ = 7px), intensity inverted and converted to 8-bit such that the full 8-bit dynamic range was utilized. Extended intensity minima were identified from this processed image (Legland, Arganda-Carreras et al. 2016) and used as markers in marker-controlled watershed segmentation of the first binary image, using the difference of Gaussian image as the intensity landscape. Watershed segmentation was restricted to the contiguous foreground-labelled regions of the first binary image with areas smaller than 2000px^2^. Areas larger than this were assumed to correspond to clusters of cells and as such yielded unreliable results from watershed segmentation. Contiguous regions of foreground-labelled pixels were identified as candidate cell objects using connected components labelling (Legland, Arganda-Carreras et al. 2016). Any cells smaller than 200px^2^ were assumed to correspond to noise, so were discarded from further analysis; those larger than or equal to 200px^2^ and smaller than 2000px^2^ were labelled as individual cells and those greater than or equal to 2000px^2^ were labelled as cell clusters. Individual cells were tracked between video frames based on their centroid coordinates using the TrackMate implementation of the Jaqaman algorithm (Jaqaman, Loerke et al. 2008, Tinevez, Perry et al. 2017). Only tracks detected for at least 5 frames were retained for analysis. Cell clusters were similarly tracked. Finally, motion characteristics for individual cells and cell clusters were calculated and recorded. These data are available at the University of Bristol data repository, data.bris, at https://doi.org/10.5523/bris.11qjhj9l4jpf2i34kob8d4eug.

### 2.7 Cytospin preparation and staining

To analyze cell morphology 5×10^4^ cells were resuspended in 100mL PBS and cytocentrifuged onto slides at 350g for 5 minutes (Thermo Scientific, Cytospin 4) before fixation in methanol for 15 minutes and staining with May-Grünwald’s (VWR Chemicals) and Giemsa (Merck Millipore) stains according to manufacturer’s instructions.

### 2.8 Analysis of reactive oxygen species

Reactive oxygen species (ROS) production was examined over time using a luminol-amplified chemiluminescence assay. 1×10^5^ macrophages were seeded in triplicate in 100μL media (HBSS, 10mM Hepes, 0.025% HSA) after which HRP (1200 U/mL) and luminol (50mM) were added at 1:2000. Following a 15-minute incubation at 37°C, macrophages were stimulated with a final concentration of 100nM phorbol 12-myristate 13-acetate (PMA) and chemiluminescence recorded for 100 minutes in 2.5-minute intervals (FLUOstar plate reader, BMG LabTech).

### 2.9 ELISA

Antibodies specific for the SARS-CoV-2 Spike protein were detected in waste apheresis cones plasma samples by ELISA using a protocol based on a previously described (Amanat, Stadlbauer et al. 2020). Plasma was heated to 56°C for 30 minutes prior to use to inactivate any potential residual virus. The SARS-CoV-2 trimeric Spike protein was produced in insect cells as previously described (Amanat, Stadlbauer et al. 2020, Stadlbauer, Amanat et al. 2020). MaxiSorp high-binding ELISA plates (NUNC) were coated with 10μg/mL Spike protein in PBS (50mL per well), and incubated overnight at 4°C. All subsequent stages of the protocol were performed at room temperature. Unbound antigen was removed with 3x washes in PBS with 0.1% Tween (0.1% PBST). Plates were blocked for 1 hour with 3% BSA/PBS to prevent non-specific binding. Plasma samples were diluted in 1% BSA/PBS and added to the coated plate (100mL per well) for 2 hours. After 3x washes with 0.1% PBST, HRP-conjugated anti-human pan immunoglobulin secondary antibody (Sigma; 1:10,000) was added (50mL per well) and incubated for 1 hour. After a final three washes in 0.1% PBST, plates were dried completely and developed with the addition of the HRP substrate OPD (SigmaFast; 100mL per well). The reaction was terminated after 30 minutes by the addition of 3M HCl (50mL per well). Optical density was measured at 492nm and 620nm on a BMG FLUOstar OMEGA MicroPlate Reader with MARS Data Analysis software. The 620nm reference wavelength measurements were subtracted from the 492nm measurement wavelengths for each well to give background corrected values. Averaged blank values from wells containing no plasma were then subtracted from all experimental values. Values were then normalized to the internal positive control. Statistical calculations were performed in Prism 8 (GraphPad) with a nonpaired t-test used to assess the difference between groups.

### 2.10 TMT labelling and high pH reversed-phase chromatography

For the surfaceome analysis, streptavidin-isolated samples were reduced (10mM TCEP, 55°C for 1h), alkylated (18.75mM iodoacetamide, room temperature for 30min.) and then digested from the beads with trypsin (2.5μg trypsin; 37°C, overnight). For the whole cell proteomics, 100μg of each sample was reduced, alkylated and digested with trypsin as above. The resulting peptides were then labeled with TMT eleven-plex reagents according to the manufacturer’s protocol (Thermo Fisher Scientific, Loughborough, LE11 5RG, UK) and the labelled samples pooled and desalted using a SepPak cartridge according to the manufacturer’s instructions (Waters, Milford, Massachusetts, USA). Eluate from the SepPak cartridge was evaporated to dryness and resuspended in buffer A (20 mM ammonium hydroxide, pH 10) prior to fractionation by high pH reversed-phase chromatography using an Ultimate 3000 liquid chromatography system (Thermo Scientific). In brief, the sample was loaded onto an XBridge BEH C18 Column (130Å, 3.5μm, 2.1mm X 150 mm, Waters, UK) in buffer A and peptides eluted with an increasing gradient of buffer B (20 mM Ammonium Hydroxide in acetonitrile, pH 10) from 0-95% over 60 minutes. The resulting fractions (4 for the surfaceome analysis and 15 for the whole cell analysis) were evaporated to dryness and resuspended in 1% formic acid prior to analysis by nano-LC MSMS using an Orbitrap Fusion Tribrid mass spectrometer (Thermo Scientific).

### 2.11 Nano-LC Mass Spectrometry

High pH RP fractions were further fractionated using an Ultimate 3000 nano-LC system in line with an Orbitrap Fusion Tribrid mass spectrometer (Thermo Scientific). In brief, peptides in 1% (vol/vol) formic acid were injected onto an Acclaim PepMap C18 nano-trap column (Thermo Scientific). After washing with 0.5% (vol/vol) acetonitrile 0.1% (vol/vol) formic acid, peptides were resolved on a 250mm × 75μm Acclaim PepMap C18 reverse phase analytical column (Thermo Scientific) over a 150 min organic gradient with a flow rate of 300nL min^-1^. Solvent A was 0.1% formic acid and Solvent B was aqueous 80% acetonitrile in 0.1% formic acid. Peptides were ionized by nano-electrospray ionization at 2.0kV using a stainless-steel emitter with an internal diameter of 30μm (Thermo Scientific) and a capillary temperature of 275°C. All spectra were acquired using an Orbitrap Fusion Tribrid mass spectrometer controlled by Xcalibur 2.1 software (Thermo Scientific) and operated in data-dependent acquisition mode using an SPS-MS3 workflow. FTMS1 spectra were collected at a resolution of 120 000, with an automatic gain control (AGC) target of 200 000 and a max injection time of 50ms. Precursors were filtered with an intensity threshold of 5000, according to charge state (to include charge states 2-7) and with monoisotopic peak determination set to peptide. Previously interrogated precursors were excluded using a dynamic window (60s +/-10ppm). The MS2 precursors were isolated with a quadrupole isolation window of 1.2m/z. ITMS2 spectra were collected with an AGC target of 10 000, max injection time of 70ms and CID collision energy of 35%.

For FTMS3 analysis, the Orbitrap was operated at 50 000 resolution with an AGC target of 50, 000 and a max injection time of 105ms. Precursors were fragmented by high energy collision dissociation (HCD) at a normalized collision energy of 60% to ensure maximal TMT reporter ion yield. Synchronous Precursor Selection (SPS) was enabled to include up to 10 MS2 fragment ions in the FTMS3 scan.

### 2.12 Proteomic Data Analysis

The raw data files were processed and quantified using Proteome Discoverer software v2.1 (Thermo Scientific) and searched against the UniProt Human database (downloaded March 2020: 165104 entries) using the SEQUEST HT algorithm. Peptide precursor mass tolerance was set at 10ppm, and MS/MS tolerance was set at 0.6Da. Search criteria included oxidation of methionine (+15.995Da), acetylation of the protein N-terminus (+42.011Da) and Methionine loss plus acetylation of the protein N-terminus (−89.03Da) as variable modifications and carbamidomethylation of cysteine (+57.021Da) and the addition of the TMT mass tag (+229.163Da) to peptide N-termini and lysine as fixed modifications. Searches were performed with full tryptic digestion and a maximum of 2 missed cleavages were allowed. The reverse database search option was enabled and all data was filtered to satisfy false discovery rate (FDR) of 5%.

Resulting protein abundance data were normalized against the total abundance of each respective sample. Subsequently, the abundance of each protein from the samples of interest was divided by the average abundance of the respective protein in the two control samples to generate Fold Change (FC) data and log-transformed to generate Log_2_FC values. As the final pre-processing step, median centering was performed on the log-transformed data. For the surfaceome dataset filtering of the total dataset to generate a cell surface-specific dataset was performed through use of the *in silico* human surfaceome database (Bausch-Fluck, Goldmann et al. 2018). Log_2_FC values were clustered with the use of Cluster 3.0 (de Hoon, Imoto et al. 2004) via average-link hierarchical clustering, and the resulting dendrograms and heatmaps were visualized with the use of Java TreeView 1.1.6r4.(Saldanha 2004) Gene Ontology (GO), KEGG pathway and network analysis of differentially expressed genes were performed through EnrichR (Kuleshov, Jones et al. 2016) and the STRING database (Jensen, Kuhn et al. 2009). The mass spectrometry proteomics data will be deposited to the ProteomeXchange Consortium via PRIDE (Perez-Riverol, Csordas et al. 2019) with the dataset identifiers PXD032801, PXD032823, and PXD032967

### 2.13 Statistical analysis

Where appropriate, statistical analysis was used to determine statistical significance and stated within the relevant figure legends. In summary, statistical analyses and generation of graphics were performed using GraphPad Prism 9 (Version 9.1.0). Details of statistical tests used and numbers of independent experiments are indicated in the respective figure legends. Standard deviation is shown where applicable. Data was confirmed to be either normally distributed (a=0.05) or non-normally distributed via the D’Agostino-Pearson omnibus test or Shapiro-Wilk test prior to further comparisons. When the data was normally distributed, an unpaired twosided t-test was used for groups of two sample types, and an ordinary one-way ANOVA test with Dunnet’s multiple comparison posthoc-test was used to compare groups of more than two sample types comparing every mean to the control mean. Alternatively, an ordinary one-way ANOVA was followed by a Tukey test to compare every mean to every other mean. For non-normally distributed data, the unpaired twosided Mann-Whitney test was used for groups of two and the Kruskal-Wallis test with Dunn’s multiple comparison posthoc-test were used to compare groups of more than two sample types comparing every mean to the control mean. For results presented as a percentage of the control, a single sample t-test and Wilcoxon test were used. Statistical significance is indicated on graphs using standard conventions, as follows: non-significant (ns), p≥0.05; *, p<0.05; **, p<0.01; ***, p<0.001; ****, p<0.0001.

## 3 Results

### 3.1 Surface marker expression on ex vivo generated macrophage subtypes

Cells from healthy donors were cultured to produce four macrophage populations; MØ control macrophages, pro-inflammatory M1 stimulated with IFNg, alternatively activated M2a macrophage stimulated with IL-4, and deactivated M2c macrophages stimulated with dexamethasone (Heideveld, Masiello et al. 2015, Gurevich, Severn et al. 2018, Hampton-O’Neil, Severn et al. 2020). Cytospins were performed on day 7 of culture to visualize distinct morphology. It was observed that classically activated M1 macrophages display lobed nuclei and cell projections in the form of filipodia, whilst the nuclei of M2c deactivated macrophages were more condensed and the cells larger (indicated by arrows, Figure 1a).

**Figure 1:**
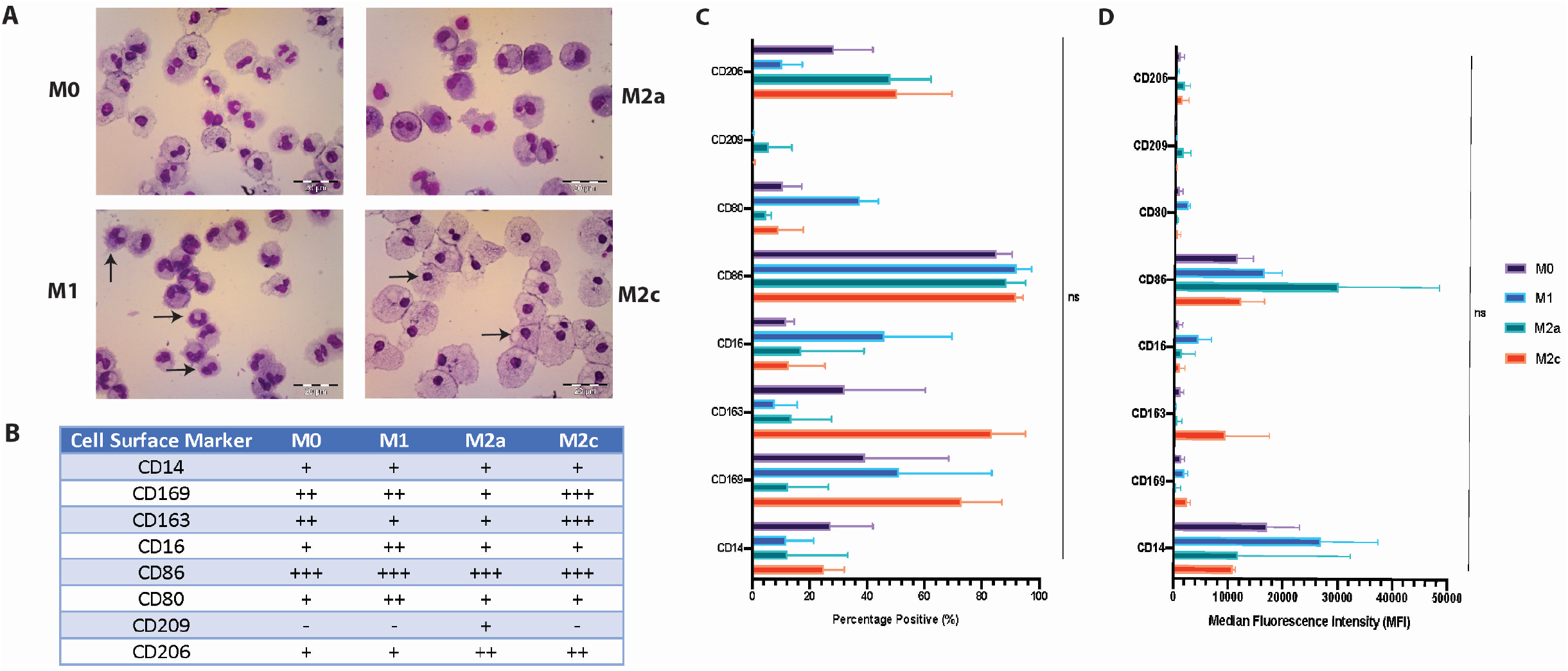
Polarized macrophage subtypes differ in cell surface marker expression and morphology in comparison to samples from patients recovered from COVID-19. A) Representative cytospins at day 7 from each of the subtypes, MØ control, M1, M2a and M2c stained with May-Grünwald’s and Giemsa’s. Scale bars 20μm, images are representative of N=3. B) Flow cytometry analysis summary of cell surface expression for key macrophage markers at day 7 of *ex vivo* culture across 4 different subtypes; CD14, CD169, CD163, CD16, CD86, CD80, CD209 and CD206. Expression presented as a scale. N=3 with a minimum of 10,000 events per sample. C) Flow cytometry analysis of cell surface expression of the following key macrophage markers at day 7 of *ex vivo* culture; CD14, CD169, CD163, CD16, CD86, CD80, CD209 and CD206. D) Median fluorescence intensity of flow cytometry markers tested in C). All flow cytometry experiments show the mean and standard deviation for the data (N=3, ns p≥0.05), with a minimum of 10,000 events per sample, significance tested using a one-way ANOVA followed by a Tukeys multiple comparison test, comparing each subtype.

Cell surface expression of the four *ex vivo* generated macrophage subtypes was explored using flow cytometry. Experiments showed that MØ unstimulated control macrophages were CD14^+^/CD169^++^/CD163^++^/CD16^+^/CD86^+++^/CD80^+^/CD209^-^ /CD206^+^, in comparison the M1 subtype were CD14^+^/CD169^++^/CD163^+^/CD16^+^/CD86^+++^/CD80^+^/CD209^-^/CD20^+^, the M2a subtype were CD14^+^/CD169^+^/CD163^+^/CD16^+^/CD86^+++^/CD80^+^/CD209^+^/CD206^++^ and finally M2c macrophages were CD14^+^/CD169^+++^/CD163^+++^/CD16+/CD86^+++^/CD80^+^/CD209^-^ /CD206^++^ (Figure 1b, c and d).

### 3.2 Functional assessment of *ex vivo*-generated macrophage subtypes

To further characterize the four macrophage populations beyond cell surface marker expression and morphology, the functionality of the cell types was also investigated. The motility of cells was assessed as an indicator of cell activity utilizing a recently described live cell imaging pipeline (Heideveld, Hampton-O’Neil et al. 2018, Hampton-O’Neil, Severn et al. 2020). Macrophages were stimulated as above utilizing MCSF-only for MØ, IFNg for M1, IL-4 for M2a and dexamethasone for M2c subtypes, without the presence of any additional chemotaxis stimuli, which have the potential to effect polarization of the cells. In this way the spontaneous motility of each macrophage subtype was assessed. Macrophages on day 7 of culture were imaged every 20 minutes, using an IncuCyte^®^ over a 12-hour time course in multiple fields of view with representative images in Supplemental Figure 1a and videos available upon publication. Figure 2a demonstrates the significantly reduced (P<0.0001) motility of M1 macrophages and increased motility of M2a macrophages when assessing the mean speed in comparison to MØ control macrophages. M2a macrophages are significantly more motile (P<0.0001), with a speed of 17.7mm in each 20-minute frame compared to 12.1mm/frame for control MØ. M2c macrophages displayed a similar mean speed to MØ control, albeit with a greater heterogeneity in the M2c population (Figure 2a). The mean Euclidean distance, a measure of distance between the start and end of a cell’s travel, from each subtype was significantly different (P<0.0001, Figure 2b). Cells in the M1 subtype travelled substantially less compared to the MØ control. M2c macrophages displayed a similar mean Euclidean distance to that of MØ control, whereas this was increased M2a in macrophages.

**Figure 2:**
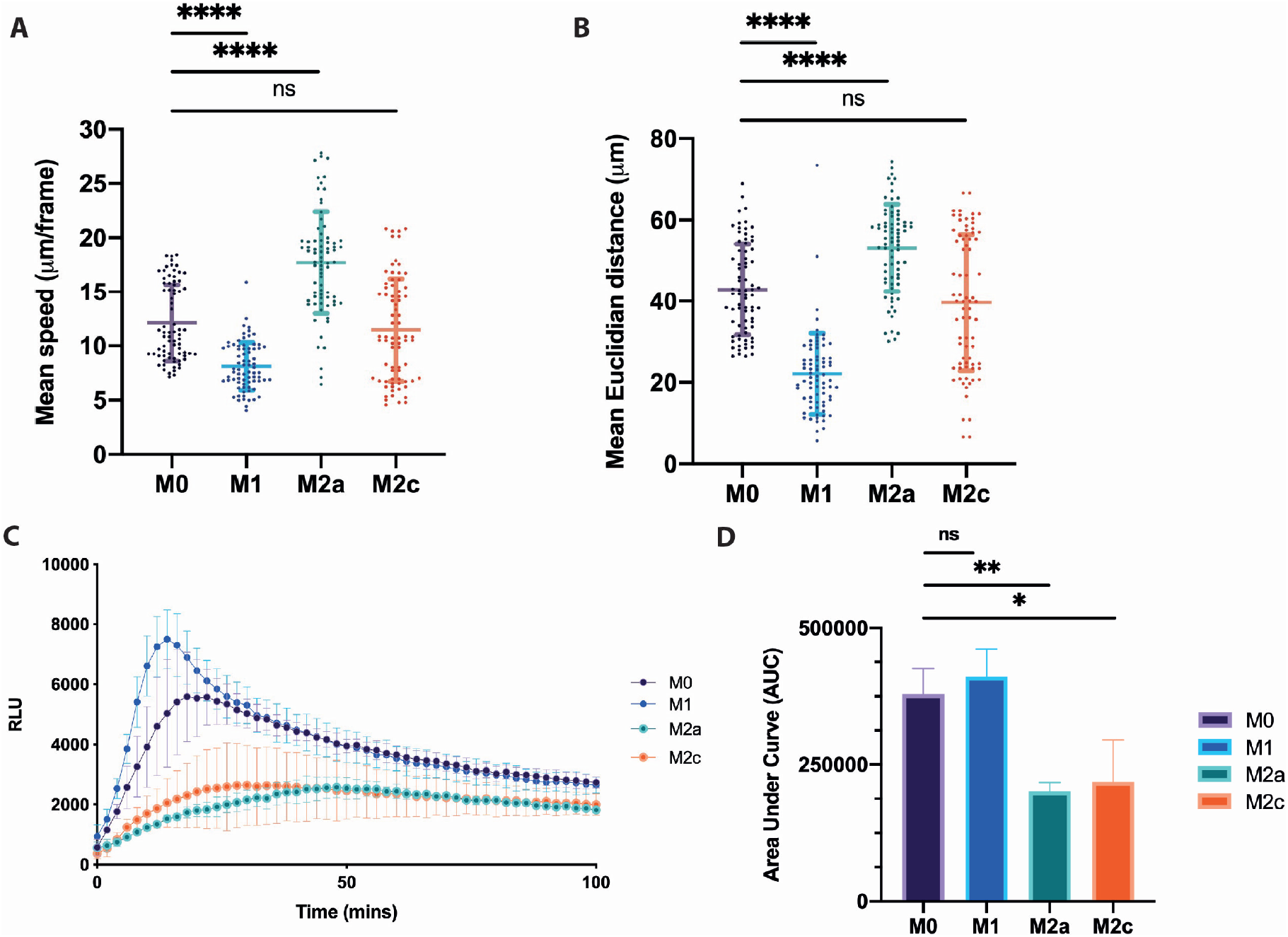
Locomotive and reactive oxygen species (ROS) analysis of ex vivo generated macrophage subtypes. A) Scatter plot with mean and standard deviation displayed of the mean speed of macrophages in each IncuCyte^®^ imaging field per 20-minute time frame of the four macrophage subtypes Kruskal-Wallis test (**** (p<0.0001)) followed by a Dunn’s multiple comparison test was performed on 75 fields of view (25 fields of view per donor), (N=3 (**** (p<0.0001)). B) Scatter plot with mean and standard deviation shown of the mean Euclidean distance macrophages cover in each imaging field per 20-minute time frame of the four macrophage subtypes. Kruskal-Wallis test (**** (p<0.0001)) followed by a Dunn’s multiple comparison test was performed on 75 fields of view (25 fields of view per donor) (N=3, **** (p<0.0001)). C) Respiratory burst formation in response to PMA stimulation measured for each of the 4 macrophage subtypes for 100 minutes. Error bars represent the standard deviation. D) Area under the curve analysis for C) where Kruskal-Wallis test (** (p<0.01)) followed by a Dunn’s multiple comparison test was performed (N=3, ** (p<0.01) and * (p<0.05)).

The production of reactive oxygen species (ROS) is a hallmark of the macrophage immune response, and thus ROS production was evaluated in each macrophage subtype. Cells were cultured and harvested on day 7 and then stimulated with 100nM phorbol 12-myristate 13-acetate (PMA) and ROS production measured using a luminol-amplified chemiluminescence assay. Each of the four subtypes generated a respiratory burst in response to PMA stimulation (Figure 2c and area under the curve analysis in Figure 2d). IFNg-induced M1 macrophages displayed the highest production of ROS as observed previously (Zhang, Choksi et al. 2013, Canton, Khezri et al. 2014); with a characteristic initial peak of ROS production observed before a plateau of continuous production. Interestingly, control MØ macrophages also exhibited ROS production levels comparable to M1 macrophages, although to a lesser degree for the initial peak, suggestive of a priming mechanism by IFNg. The M2a and M2c subtypes generated ROS at significantly reduced rates to MØ macrophages (P = <0.01 and P = 0.05 respectively).

### 3.3 Whole-cell quantitative proteomics reveal the extended scale of macrophage subtype plasticity

We next pursued a global overview of the phenotype continuum of macrophage plasticity through a tandem mass tagging (TMT)-based whole-cell proteomics approach comparing polarity states simultaneously. MØ control, M1, M2a and M2c lysates produced from the same donor material, were TMT-labelled before proceeding on to mass spectrometric and computational analysis (Figure 3a). Figure 3b displays the log_2_ fold change (log_2_ FC) of protein levels in individual samples relative to the originating MØ control population, demonstrating the broad variation between subtypes in the dataset as a whole, comprising a total of 5435 proteins (Supplemental Table 1). Sub-cluster analysis for the heatmap in Figure 3b highlights key broad differences between the macrophage subtypes in the group of differentially expressed proteins. Broad patterns of specific biological regulation are immediately identifiable amongst this cluster-based analysis, such as a strong interferon/cytokine response in M1 macrophages and the upregulation of cell-matrix adhesion and collagen fibrils in M2c macrophages. A filtered dataset of differentially expressed proteins (absolute mean log_2_ FC>1) is provided in Supplemental Table 2.

**Figure 3:**
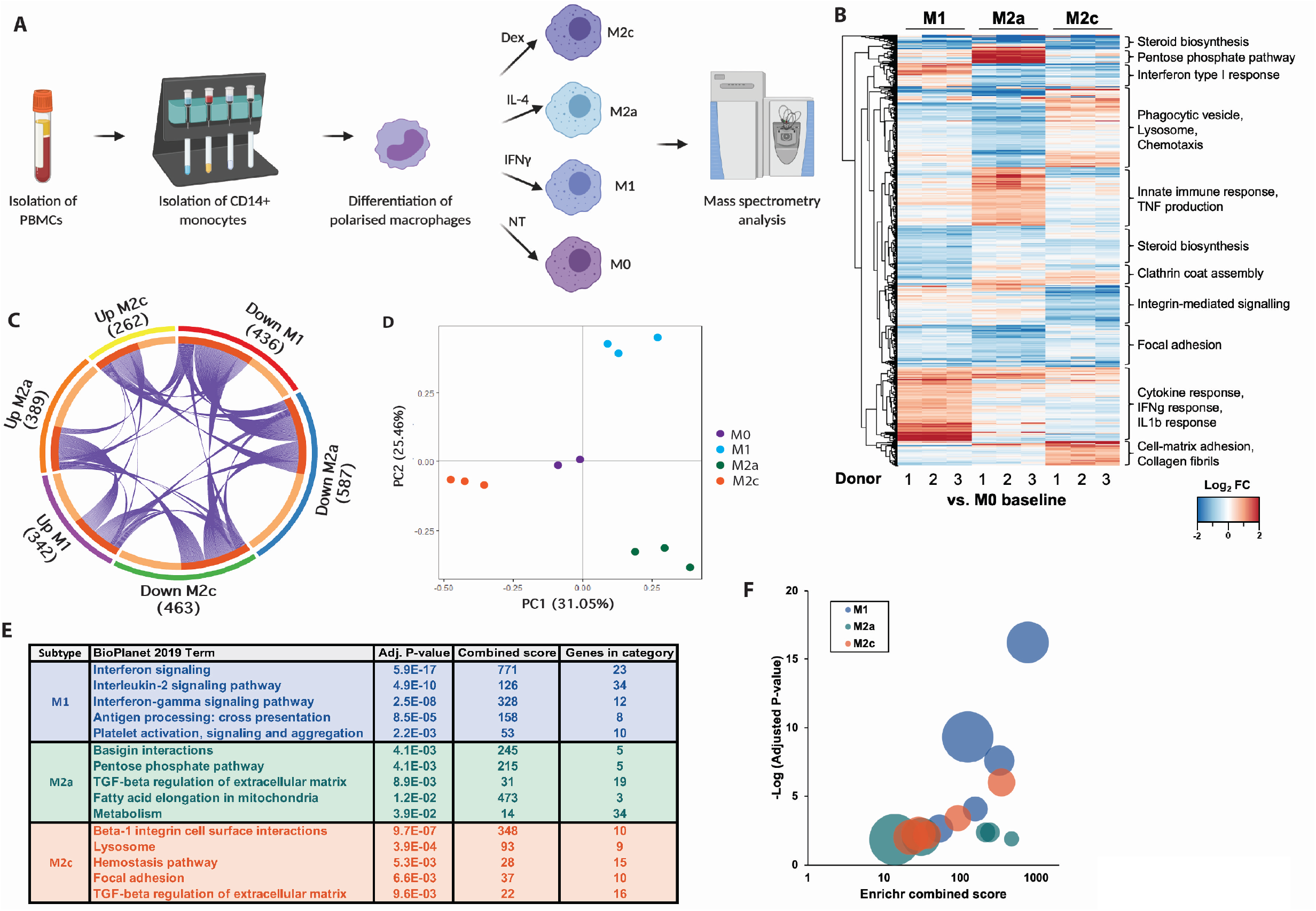
Semi-quantitative TMT proteomics elucidate the protein changes underlying macrophage polarization. A) Experimental design for tandem mass tagged (TMT) proteomic analysis, CD14^+^ cells were isolated from donated human peripheral blood before being stimulated to M1, M2a, M2c or MØ control subtypes. B) Heatmap visualization of the TMT proteomic analysis generated from log_2_ fold changes in expression of whole-cell lysates with subcluster descriptions. The figure legend denotes lower expression are represented in blue whilst red higher expression, FDR set at 5%. Log_2_FC values were clustered with the use of Cluster 3.0 (de Hoon, Imoto et al. 2004) via average-link hierarchical clustering, and the resulting dendrograms and heatmaps generated. C) Chord plot summarizing the proportion of up and downregulated proteins for each of the M1, M2a and M2c macrophage subtypes with values representing this for each of the subtypes. D) Principal component analysis of the four macrophage subtypes from the total proteomic dataset. E) Top 5 enriched meta-categories per macrophage subtype per the Enrichr BioPlanet 2019 enrichment module. F) Bubble plot from results in E.

These proteins were then processed through the STRING database creating protein-protein interaction networks to facilitate visualization and analysis (Supplemental Figures 2a, b and c). Further analysis showed that the abundance of 2552 proteins remained fully consistent (absolute mean log_2_ FC >-1 and <1) across all polarized subtypes and MØ control, many of these comprising housekeeping genes (Supplemental Table 3). This observation is consistent with the common origin of all subtypes from the same macrophage population, but it is important to note that all subtypes differ significantly in >50% of the proteome. The relationships between the 3 polarized macrophage subtypes can be visualized in the chord plot (Figure 3c), differentially expressed proteins and their directionality are clustered by co-occurrence between subtypes. As an additional confirmation of these differential patterns of gene expression regulation, we performed a principal component analysis which identified four distinct and easily separable clusters, each corresponding to a macrophage subtype with the MØ control central to this analysis (Figure 3d).

Finally, a subset-centered investigation of differentially expressed proteins, based on gene set enrichment analysis (GSEA) and performed with the Enrichr BioPlanet 2019 module, confirmed that M1 macrophages exhibited a heavily pro-inflammatory profile, as anticipated, but also that M2a were skewed towards chemokine-based signaling and M2c possessed a broad regulatory and extracellular matrix-centric profile. A summary table displaying the top 5 enriched meta-categories for each macrophage subtype compared to MØ is presented in Figure 3e, with a corresponding bubble plot for ease of visualization in Figure 3f (where bubble size denotes the number of genes in the category shown in the table), and the complete enrichment results are provided in Supplemental tables 4 – 6. More specific and biologically-relevant alterations in protein abundance included the M1 upregulation of Mx1 and Mx2, (Aebi, Fah et al. 1989) two IFNg-induced GTP-binding proteins with anti-viral activity, as well as increased abundance of the anti-viral enzymes OAS1, 2 and 3 (Eskildsen, Justesen et al. 2003). The M2a subtype-upregulated proteins included CLEC4a (Bates, Fournier et al. 1999) 5A,(Bakker, Baker et al. 1999), 10A (Higashi, Fujioka et al. 2002) and 16A (Zouk, D’Hennezel et al. 2014), the C-type lectin family members which mediate immune activity (Geijtenbeek and Gringhuis 2009); platelet factor 4 a chemotactic agent released during platelet aggregation (Deuel, Senior et al. 1981); and integrin alpha-M (Owen, Campbell et al. 1992), involved in adhesive interactions of monocytes and granulocytes. Conversely, M2c broadly upregulated genes from the collagen family, which in turn were downregulated in the M1 subtype. M2c also upregulated integrin alpha-V, responsible for the recognition of the RGD sequence in extracellular matrix constituents (Weinacker, Chen et al. 1994, Kapp, Rechenmacher et al. 2017).

### 3.4 Quantitative “surfaceome” analysis of ex vivo-generated macrophage subtypes

Consecutive surface biotinylation and streptavidin pull downs were performed to facilitate identification of cell surface proteins across the four cultured macrophage populations, before TMT-based proteomic analysis. Differential protein expression was evaluated and visualized by comparison with MØ control macrophages (Supplemental Figure 3a and Supplemental Table 7). These datasets were filtered to specifically identify cell surface proteins, using the Cell Surface Protein Atlas (CSPA), removing confounding intracellular proteins (arising due to unspecific association with biotinylated surface proteins (Bausch-Fluck, Goldmann et al. 2018)). The resultant dataset and heatmap shown in Figure 4a (Supplemental Table 8) were thus curated to 90 proteins from an original total of 921, and all subsequent analyses were performed with this dataset. A new cluster-based analysis was performed on this dataset, still demonstrating broad differences in gene expression between the subtypes with identifiable patterns, albeit to a lesser degree than the whole-cell proteome data (Figure 4a).

**Figure 4:**
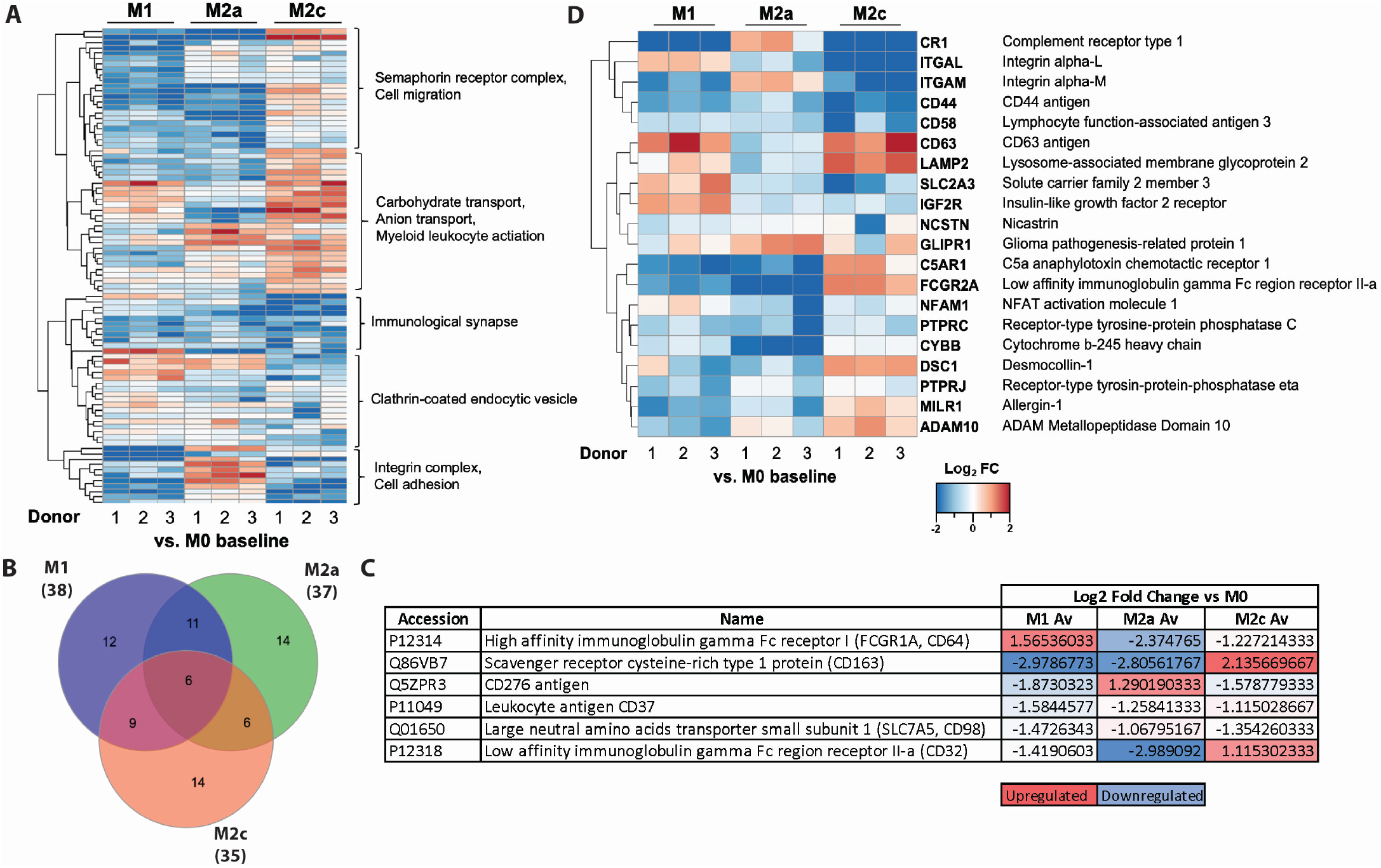
The macrophage “surfaceome” is dynamically altered upon polarization. A) Heatmap visualization of the tandem mass tagged (TMT) surfaceome analysis filtered for surface proteins using the *in silico* database (Bausch-Fluck, Goldmann et al. 2018) (accessed June 2020) generated from Log_2_ fold changes in expression in M1, M2a and M2c subtypes with the MØ control as a baseline. Descriptions of the subclusters are also included. Lower expression represented in blue whilst red denotes higher expression. Log_2_FC values were clustered with the use of Cluster 3.0 (de Hoon, Imoto et al. 2004) via average-link hierarchical clustering, and the resulting dendrograms and heatmaps generated. B) Venn diagram of the consistently up and down regulated (log_2_ fold change ≤ 1 and ≥ 1) proteins between three subtypes using the MØ control macrophages as a baseline for comparison. C) Table illustrating results from Figure 4b, presented are the proteins either significantly up- or downregulated for each of the three subtypes when expressed as a Log_2_ fold change (log_2_ fold change ≤ 1 and ≥ 11) against the MØ control D) Heatmap visualization of the “myeloid cell activation involved in immune response” GO category term in the surface filtered TMT dataset. N=3 for macrophage subtypes and N=2 for MØ control, FDR was set at 5%.

A summary of differential expression of up and downregulated hits between the subtypes is illustrated in Figure 4b. These differentially expressed proteins (*vs*. control MØ) totaled 35 proteins in M2c macrophages, compared to 38 and 37 proteins for the M1 and M2a subtypes respectively, whilst only 6 were identified as overlapping between the subtypes (Figure 4b and c). These data also highlight key differences between M2a and M2c subtypes where these two subtypes shared the least commonly up or down regulated proteins. Significantly up or downregulated proteins shared between all 3 subtypes (*vs*. control MØ) are displayed in Figure 4c. STRING analysis was also performed for up and downregulated hits (Supplemental Figures 3b-g). Evaluation of the Gene Ontology (GO term) category “myeloid cell activation in immune response” showed that M2c displayed the broadest response compared to control MØ, significantly upregulating 5 out of 20 hits, namely: CD63 TIMP1 receptor, lysosome associated membrane glycoprotein 2, low affinity immunoglobulin gamma FC region receptor II, desmocollin-1, allegrin-1, and ADAM 10 (Figure 4d and Table 1).

**Table 1:** Surfaceome analysis using Gene Ontology (GO term) category “myeloid cell activation in immune response”.

| Uniprot ID | Gene symbol | Extended gene name | Log<sub>2</sub>FC<br>M1/M0<br>(Avg.<br>3) | Log<sub>2</sub>FC<br>M2a/M0<br>(Avg. 3) | Log<sub>2</sub>FC<br>M2c/M0<br>(Avg. 3) |
| --- | --- | --- | --- | --- | --- |
| P17927 | CR1 | Complement receptor type 1 | 3.02 | 0.74 | 3.36 |
| P20701 | ITGAL | Integrin alpha-L | 0.66 | 0.84 | 4.02 |
| P11169 | SLC2A3 | Solute carrier family 2, facilitated glucose transporter member 3 | 0.89 | 0.61 | 1.42 |
| P16070 | CD44 | CD44 antigen | 1.31 | 0.86 | 1.78 |
| P19256 | CD58 | Lymphocyte function-associated antigen 3 | 0.59 | 0.73 | 1.43 |
| P11215 | ITGAM | Integrin alpha-M | 1.51 | 0.69 | 2.26 |
| P11717 | IGF2R | Cation-independent mannose-6-phosphate receptor | 1.04 | 0.48 | 0.58 |
| P08575 | PTPRC | Receptor-type tyrosine-protein phosphatase C | 0.79 | 1.35 | 0.56 |
| Q8NET5 | NFAM1 | NFAT activation molecule 1 | 0.25 | 0.77 | 0.28 |
| P04839 | CYBB | Cytochrome b-245 heavy chain | 0.49 | 0.91 | 0.07 |
| Q12913 | PTPRJ | Receptor-type tyrosine-protein phosphatase eta | 1.02 | 0.34 | 0.18 |
| Q92542 | NCSTN | Nicastrin | 0.46 | -0.03 | 0.67 |
| P48060 | GLIPR1 | Glioma pathogenesis-related protein 1 | 0.38 | 1.08 | 0.68 |
| Q7Z6M3 | MILR1 | Allergen-1 | 0.52 | 0.85 | 0.50 |
| O14672 | ADAM10 | Disintegrin and metalloproteinase domain-containing protein 10 | 0.17 | 0.40 | 0.80 |
| Q08554 | DSC1 | Desmocollin-1 | 0.98 | 0.91 | 1.06 |
| P21730 | C5AR1 | C5a anaphylatoxin chemotactic receptor 1 | 1.83 | 2.05 | 0.79 |
| P12318 | FCGR2A | Low affinity immunoglobulin gamma Fc region receptor II-a | 1.42 | 0.40 | 1.12 |
| P08962 | CD63 | CD63 antigen | 1.44 | -0.27 | 1.48 |
| P13473 | LAMP2 | Lysosome-associated membrane glycoprotein 2 | 0.37 | 0.42 | 1.43 |

### 3.5 Dexamethasone treatment of pro-inflammatory M1 macrophages shifts cell polarity towards a de-activated M2 phenotype

Dexamethasone has been widely used as a treatment for a variety of immunopathologies since the 1960’s. More recently dexamethasone has been used in the treatment of hospitalized COVID-19 patients, where M1 macrophages likely contribute to the cytokine storm exhibited in respiratory disorders (Group, Horby et al. 2020). We therefore utilized this model system to investigate the effect of dexamethasone on M1 macrophages. Isolated CD14^+^ monocytes were polarized to M1 macrophages using IFNg before simultaneous treatment with dexamethasone for 48 hours to mimic disease pathology in patients, analysis was then conducted on day 7. Dexamethasone-treated M1 macrophages had increased levels of CD163 and CD206, albeit neither were found to be significantly increased; both of which are known markers of M2 macrophages (Gundra, Girgis et al. 2014, Heideveld, Hampton-O’Neil et al. 2018). Conversely, the M1 markers CD80 and CD86 remained unchanged with dexamethasone treatment (Figure 5a and Supplemental Figure 4a). Interestingly, an accompanying increase in TNFαR and ACE2 (mean = 29.19% vs. 1.67% and 27.93% vs. 2.88%, for control and dexamethasone treatment respectively) surface expression was observed, albeit insignificant due to donor variation and low MFI expression (Figure 5a and b and Supplemental Figures 4a and b). Treatment with dexamethasone for 24-hours displayed a similar, albeit weaker trend than that of the 48-hour treatment (data not shown). In addition, the effect of dexamethasone treatment on capacity for ROS generation was tested, a 48-hour treatment did not induce a reduction in ROS levels, when compared to untreated M1 macrophages (Supplemental Figure 4e).

**Figure 5:**
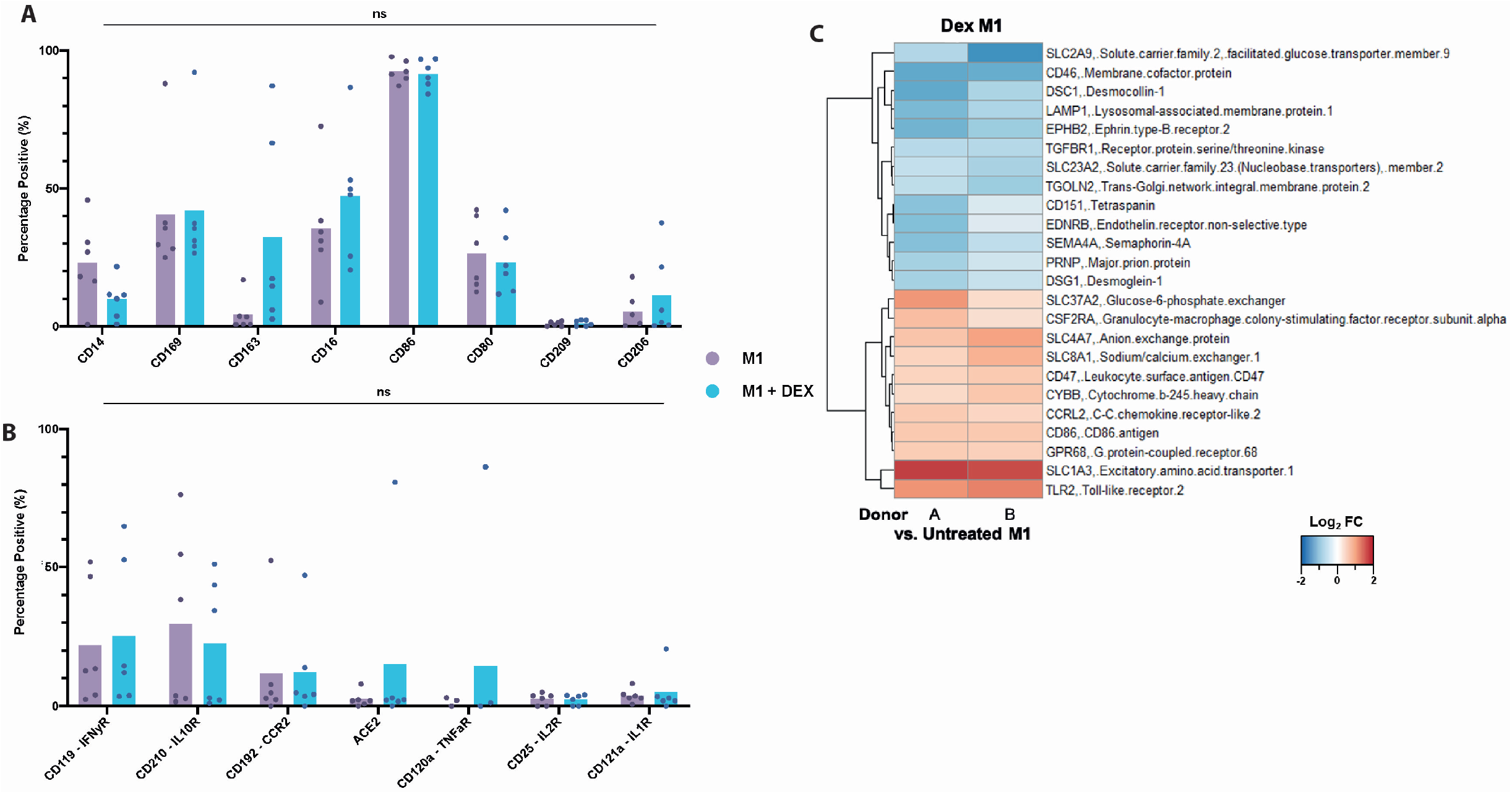
Dexamethasone treatment of M1 macrophages induces partial polarization towards the M2c phenotype. A and B) Flow cytometry analysis of dexamethasone treatment for 48 hours on M1 induced macrophages to determine plasticity of inflammatory macrophages. Percentage positive of population shown. Bar shows mean with each point representing individual samples (N=6, ns p>0.05), with a minimum of 10,000 events per sample, significance tested using a two-tailed t-test. A) Flow cytometry analysis included subtype specific markers in addition to markers of the inflammatory response as including: CD14, CD169, CD163, CD16, CD86, CD80, CD209 and CD206. B) Analysis included markers synonymous with COVID-19 infection and inflammation: CD119, CD210, CD192, ACE2, CD120a and CD121a. C) Heatmap visualization of proteomic comparison showing significantly up or downregulated proteins when comparing dexamethasone treated samples against M1 as a baseline.

To further investigate the effect of dexamethasone; MØ control, M1, M2c, and M1 dexamethasone-treated cells were surface biotinylated and TMT-labelled in a semi-quantitative proteomic approach. Results were again filtered using the CSPA database, resulting in a final dataset of 160 cell surface proteins (Supplemental Table 9). Supplemental Figure 4c gives a broad comparison of the effects of dexamethasone displayed as log_2_ FC, using MØ control as a baseline. Interestingly, only 24 of the 160 surface proteins detected were significantly differentially expressed on comparison of untreated and dexamethasone treated M1 cells (Figure 5c, Supplemental Figure 4d and Table 2). For example, SLC2A9, CD46 and LAMP-1 were significantly decreased whereas SLCA3, TLR2 and CSF2RA were significantly increased upon treatment with dexamethasone, indicating that a 48-hour treatment only partially affects the surface protein abundance of these cells.

**Table 2:** Significantly up- or down-regulated hits from the surfaceome analysis of dexamethasone treatment, expressed as a Log_2_ fold change against M1 samples.

| Uniprot ID | Gene symbol | Extended gene name | Log<sub>2</sub>FC<br>DexM1-A/M1 | Log<sub>2</sub>FC<br>DexM1-B/M1 |
| --- | --- | --- | --- | --- |
| P43003 | SLC1A3 | Excitatory amino acid transporter 1 | 1.79 | 1.71 |
| O60603 | TLR2 | Toll-like receptor 2 | 1.11 | 1.26 |
| B5M450 | SLC4A7 | Anion exchange protein | 0.64 | 0.97 |
| Q8TED4 | SLC37A2 | Glucose-6-phosphate exchanger | 1.08 | 0.36 |
| P32418 | SLC8A1 | Sodium/calcium exchanger 1 | 0.47 | 0.83 |
| P42081 | CD86 | CD86 antigen | 0.59 | 0.61 |
| A0A024R6D5 | GPR68 | G protein-coupled receptor 68 | 0.56 | 0.53 |
| Q08722 | CD47 | Leukocyte surface antigen CD47 | 0.49 | 0.58 |
| P15509 | CSF2RA | Granulocyte-macrophage colony-stimulating factor receptor subunit alpha | 0.71 | 0.31 |
| P04839 | CYBB | Cytochrome b-245 heavy chain | 0.39 | 0.62 |
| O00421 | CCRL2 | C-C chemokine receptor-like 2 | 0.57 | 0.44 |
| O75942 | PRNP | Major prion protein | -0.71 | -0.36 |
| Q02413 | DSG1 | Desmoglein-1 | -0.70 | -0.45 |
| Q5T7S2 | TGFBR1 | Receptor protein serine/threonine kinase | -0.56 | -0.60 |
| A0A024RCB3 | CD151 | Tetraspanin | -0.93 | -0.23 |
| D3DVZ8 | SLC23A2 | Solute carrier family 23 (Nucleobase transporters) member 2 | -0.48 | -0.69 |
| P24530 | EDNRB | Endothelin receptor non-selective type | -1.00 | -0.21 |
| A0A5H1ZRP2 | TGOLN2 | Trans-Golgi network integral membrane protein 2 | -0.52 | -0.78 |
| Q9H3S1 | SEMA4A | Semaphorin-4A | -0.96 | -0.52 |
| A0A024RDY3 | LAMP1 | Lysosomal-associated membrane protein 1 | -1.05 | -0.65 |
| P29323 | EPHB2 | Ephrin type-B receptor 2 | -1.13 | -0.77 |
| Q08554 | DSC1 | Desmocollin-1 | -1.24 | -0.66 |
| Q9NRM0 | SLC2A9 | Solute carrier family 2 facilitated glucose transporter member 9 | -0.63 | -1.51 |
| P15529 | CD46 | Membrane cofactor protein | -1.25 | -1.19 |

### 3.6 Investigation of COVID-19 related surface marker expression on ex vivo macrophage subtypes

We also tested multiple surface markers by flow cytometry that are highlighted as clinically relevant in the COVID-19 literature, including; IFNg receptor (IFNgR, CD119), interleukin-10 receptor (IL-10R, CD210), C-C chemokine receptor 2 (CCR2, CD192), tumor necrosis factor-α receptor (TNFαR), interleukin-1 receptor (IL-1R, CD121a), and angiotensin converting enzyme 2 (ACE2), the SARS-CoV-2 receptor (Huang, Wang et al. 2020, McGonagle, Sharif et al. 2020, Mehta, McAuley et al. 2020, Moore and June 2020, Zhou 2020). Expression of IFNgR and IL-10R was high and remained consistent whereas TNFαR expression was minimal and ACE2 prevalence remained modest across *ex vivo* cultures. Interestingly, M2c macrophages demonstrated elevated expression of CCR2, and to a lesser extent IL-1R (Supplemental Figures 5c and d). Detection was also attempted for interleukin-6 receptor (IL-6R), C-X-C motif chemokine receptor 3 (CXCR3) and interleukin-2 receptor (IL-2R), but expression was very low across the subtypes (data not shown).

### 3.7 Response to COVID-19 infection by the CD14^+^ monocyte/macrophage population

We utilized this workflow for the phenotypic assessment of macrophages to investigate cells within a disease setting. The effect of COVID-19 on monocyte populations was assessed in patients recovering from COVID-19. In order to circumvent the potential re-polarization of populations through extended culture, the CD14^+^ monocyte populations isolated from anonymous convalescent plasma (CP) apheresis waste samples were compared to standard apheresis donation waste samples as a control. The presence of antibodies against the SARS-Cov-2 Spike protein were measured using an ELISA assay confirming COVID-19 infection in the CP samples and excluding asymptomatic individuals from controls (P<0.0001, Figure 6a). The yield of CD14^+^ monocytes obtained through CD14^+^ microbead isolation was significantly reduced in CP samples compared to controls (P<0.05, Figure 6b), where a mean of 4.67×10^8^ PBMCs isolated from control samples yielded a mean of 9.62×10^7^ CD14^+^ monocytes whilst in CP samples a mean of 2.26×10^8^ PBMCS were obtained and from this only 8.92×10^6^ CD14^+^ cells were present. There were no differences in cell size or granularity between control and CP samples when assessed using FSC/SSC; when scatter quadrants were quantified insignificant differences between control and CP samples were found (Supplemental Figures 5a and b). When expression of cell surface markers was analyzed immediately following isolation, we observed a significant decrease (P<0.05) in CD80 surface expression, a marker of M1 macrophages for CP samples (mean = 8.5%±5.8 of population) compared to controls (mean = 25.8%±15.9 of population). Expression of certain markers, such as the M2c surface marker CD163 was increased albeit not significantly, while other key M2c markers such as CD169 remained consistent regardless of whether COVID-19 infection had occurred (Figure 6c, d and e, ns, P≥0.05). Taken together, the combined reduction in both CD14^+^ cells and CD80 expression suggests a specific depletion in M1 classically activated macrophage population in samples from individuals recovering from COVID-19 infection.

**Figure 6:**
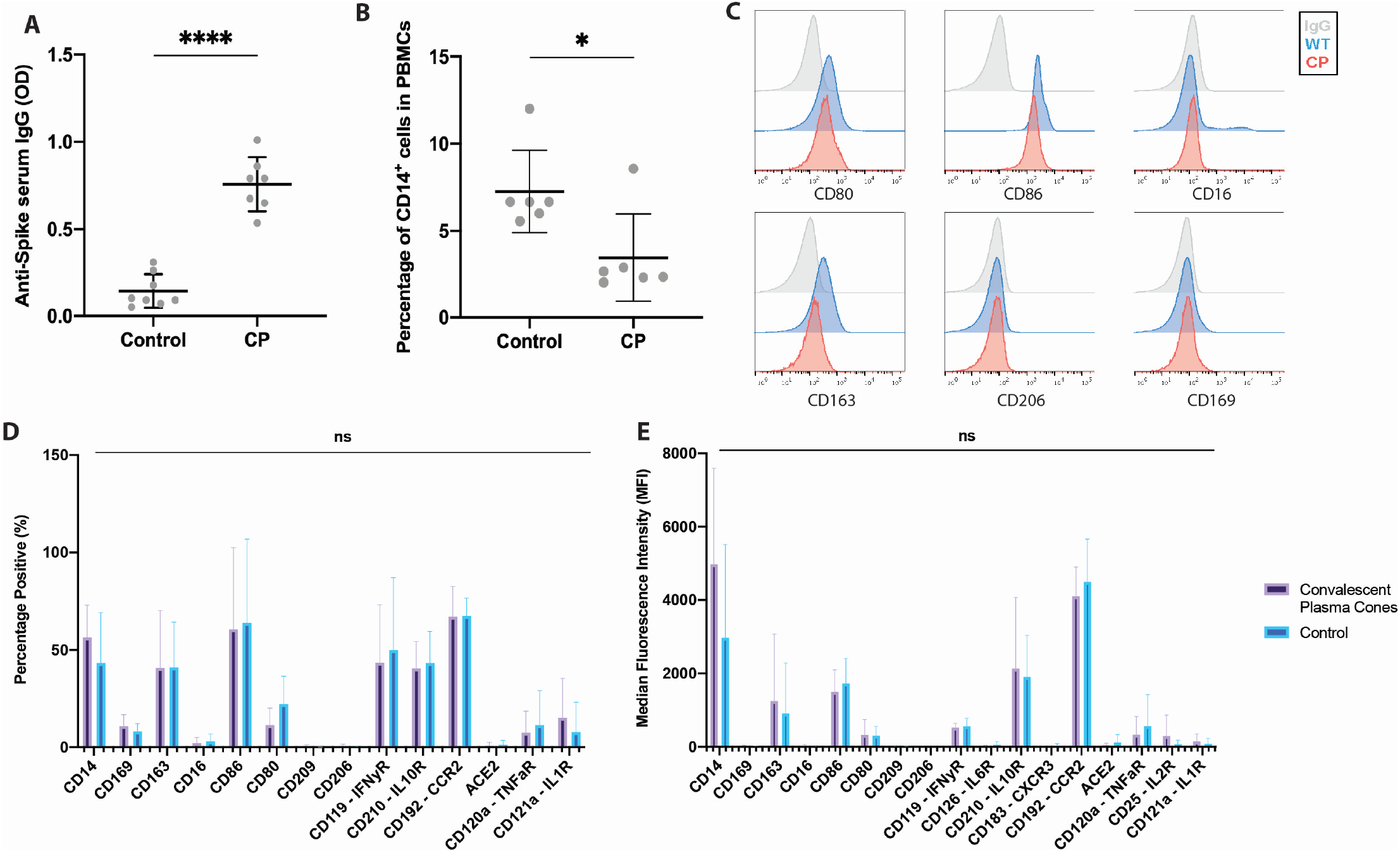
Effect of prior COVID-19 infection on *ex vivo* macrophage populations. A) Analysis of the presence of antibodies towards the Spike protein from COVID-19 infection in convalescent plasma samples (CP) and controls using an enzyme linked immunosorbent assay (ELISA). Graph shows the mean and standard deviation for the data (control N=8, CP N=7, **** p<0.0001), significance tested using a two-tailed unpaired t-test. B) Yield of CD14^+^ cells isolated as a percentage from PBMNCs is significantly reduced for convalescent plasma cones when compared to control samples. Scatter plot with the mean and standard deviation of the sample group ((N=6, * (p<0.05), significance tested using a two-tailed unpaired t-test. C) Representative histograms of flow cytometry analysis of the CD14^+^ isolated cells immediately after isolation using key macrophage markers, IgG control in grey, control in blue and CP samples in red. Histograms are representative of N=6 with a minimum of 10,000 events per sample. D) Surface markers used to investigate macrophage specific markers immediately after isolation for control and CP samples, including: CD14, CD169, CD163, CD16, CD86, CD80, CD209, CD206, CD119, CD210, CD192, ACE2, CD120a and CD121a. E) Median fluorescence intensity of flow cytometry markers tested in C). All flow cytometry experiments show the mean and standard deviation for the data (N=9, ns p≥0.05), with a minimum of 10,000 events per sample, significance tested using a two-tailed Mann-Whitney test.

## 4 Discussion

This study has extensively characterized an *ex vivo* macrophage model that produces four distinct primary macrophage subtypes. Our proteomic data and functional analysis, conducted on macrophage subtypes produced from the same donor material, confirm that substantial differences exist between the cell types induced by polarization changes. Importantly, using a whole-cell semi-quantitative TMT proteomic approach, we demonstrate distinct differential expression of proteins between the subtypes, corresponding to macrophage phenotype and function. Within the proteome dataset the majority (>50%) of proteins exhibit differential expression in abundance between the subtypes, highlighting the distinct cellular changes macrophages undergo during polarization. We identify specific profiles for M1 pro-inflammatory subtype and distinguish between M2a and M2c subtypes, which skew to chemokine or ECM-based proteins respectively. We identify a cluster of 6 surface proteins, using surfaceome studies, that are shared and consistently up or downregulated across 3 subtypes (M1, M2a and M2c) when compared to the MØ control (Figure 4b and c). Notably, key differences between M2a and M2c macrophages are distinguished within this dataset, where M2a and M2c exhibited the least commonly up or down regulated proteins.

Interestingly, our dataset show that macrophages exist beyond the dichotomy of the M1/M2 subtypes, indicated by the expression of pro-inflammatory markers in the M2c macrophages surface proteome. For example, complement receptor I (CD35) and Integrin alpha-M (CD11b) were increased in M1 and M2c cells compared to control or M2a subtypes, thus M2c cells appear to be as activated or primed to activate as M1 macrophages. Within the Gene Ontology category term “myeloid cell activation involved in immune response” the M2c subtype displayed the most prominent response compared to control, significantly upregulating 5 out of 20 hits, further supporting the pro-inflammatory priming of this subtype.

Furthermore, dexamethasone treatment of M1 macrophages, strongly induced expression of M2c marker CD163 but did not coincide with a reduction in the M1 marker CD80 expression. In functional assays treatment of M1 macrophages with dexamethasone for 48 hours did not significantly alter the ROS production capabilities of these cells, which exhibited a respiratory burst in line with an M1 phenotype rather than that observed in M2c macrophages. These intermediate or overlapping features of macrophage polarity have also been observed by others, although previous focus has been on transcriptomics and hypoxia using a simple M1/M2 model (Xue, Schmidt et al. 2014, Raggi, Pelassa et al. 2017). These data suggest that macrophage polarization is a transient continuum, with cells able to move along this sliding scale when necessary. It is important to note that a large proportion of the proteome, 2552 of the 5435 proteins identified within the whole cell dataset, remains static between all four macrophage subtypes, providing an explanation for how macrophages enable swift polarity changes along a continuum by maintaining a core set of proteins across all subtypes whose protein expression remain constant.

A comparison of the differentially expressed proteins detected in this study with those identified by Court *et al* (Court, Petre et al. 2017), displayed extremely high concordance between both datasets when investigating the three subsets, M1 (23 differentially expressed proteins matching direction out of 27 co-occurring in both lists), M2a (31/31) and M2c (18/23). These results reinforce the expanded scope of the dataset in this study and provide additional confirmation that the differential expression between the subtypes is reproducible and biologically distinct. Furthermore, our dataset is richer, with over 4-fold the number of detected differentially expressed proteins. As often observed in proteomic comparisons, we were unable to detect approximately 25% of the proteins reported by Court *et al*., which may also reflect differences in the peptide databases used. The increased scope of the integrative proteomic analyses performed in our study, reinforced by broad concordance with previous results from Court *et al*. (Court, Petre et al. 2017), provide additional confirmation and establish this study as an unparalleled investigation of macrophage subtypes and polarity *ex vivo*.

The functional studies discerned specific differences between the subtypes. Live cell imaging identified motility differences, with M1 macrophages being significantly less motile than control MØ or M2 subtypes, perhaps because they are potentially awaiting a signal for further activation. The significantly increased speed and distance travelled observed in M2a macrophages in comparison to MØ corroborates findings that alternatively activated macrophages are more motile than control macrophages, whilst M1 macrophages display significantly less motility (Hind, Lurier et al. 2016). Therefore, motility is significantly impacted by the polarization state of macrophages and can be utilized as an identifying feature of each subtype. Furthermore, when dissecting the ROS response between subtypes, we observed significant differences in ROS production between different macrophage subtypes. The response of MØ control macrophages was comparable to M1 macrophages, both of which had significantly higher ROS production compared to M2a or M2c. Therefore, ROS production is linked to the polarization state of macrophages and, further still, reflects the position of macrophages on the polarization continuum. In our study, we did not observe an alteration in ROS generation when M1 macrophages were treated with dexamethasone, whereas other reports suggested that dexamethasone increases ROS activity in M2 macrophages upon a 48-hour or 7-day treatment, perhaps due to conversion to the M2c subtype (Kraaij, van der Kooij et al. 2011). Suggesting additional changes in cell polarity are required in dexamethasone treated cells to reach a polarization state where there is a direct impact on ROS machinery. This raises the question as to whether M-CSF only stimulated macrophages are appropriate for use as a truly unpolarized macrophage in contrast to specific subtypes, when investigating the effects of macrophages *ex vivo* as a model system. Whilst seminal work investigating nursing behavior of macrophages used M-CSF only stimulated macrophages (Chow, Huggins et al. 2013, Ramos, Casu et al. 2013), more recent work has advanced our knowledge further by specifically utilizing dexamethasone alongside M-CSF to induce the M2c nurturing macrophages (Heideveld, Masiello et al. 2015, Heideveld, Hampton-O’Neil et al. 2018, Hampton-O’Neil, Severn et al. 2020). Therefore, the MØ control in the case of the work described in this manuscript is acceptable as it is compared to additionally stimulated macrophages.

To understand the effects of dexamethasone treatment used for COVID-19 patients (Group, Horby et al. 2020) on macrophage polarity, we sought to confirm whether simultaneous exposure to IFNg and dexamethasone causes a shift from the M1 phenotype to a M2c phenotype. We observed that a 48-hour dexamethasone treatment induced increased expression of CD163, through both proteomic and flow cytometry analysis. However, surface markers distinguishing between M1 and M2c such as CD169 and CD80 did not change. The glucocorticoid methylprednisolone induced M2c polarization in mouse studies, with a marked stimulation of CD163 expression and an accompanying decrease in M1 macrophage levels (Tu, Shi et al. 2017). Taken together, these observations suggest that use of dexamethasone in patients likely polarizes M1 macrophages towards M2c, thus interrupting the cytokine storm frequently reported in severe COVID-19 patients (Huang, Wang et al. 2020, Mehta, McAuley et al. 2020, Moore and June 2020). A dexamethasone treatment period of 48-hours was chosen for this study, but it is plausible that longer exposure could lead to broader and more prolonged alterations in the primary care setting. Alternatively, *ex vivo* conditions used here may not permit the generation of a fully plastic phenotype.

Finally, using CP samples from patients recovered from COVID-19 infection we demonstrate the retention of a COVID-19 footprint on the macrophage population beyond the 28 days recovery period. We were unable to confirm the observation by Zhang *et al*. of an altered macrophage cell size in peripheral blood of COVID-19 infected individuals when analyzed using flow cytometry (Zhang, Guo et al. 2020), likely due to differing sample types or patient disease severity. There is a significant decrease in expression of the M1 marker CD80 and a significant reduction in yield for both PBMNCs and CD14^+^ cells in CP samples compared to controls, correlating with a decrease in the monocyte population and particularly M1 macrophages as a consequence of viral infection (Sang, Miller et al. 2015). Therefore illustrating functional adaptivity and a continuum of plasticity based upon environmental cues *in vivo* (Stout and Suttles 2005). This suggests activation and subsequent apoptosis of the M1 population during and following infection; however further work is required to confirm this. Unfortunately, as the samples were acquired anonymously, we were unable to access hospitalization status or evaluate whether the patients had been treated with dexamethasone. However, sample data accessed for a different study (Mankelow, Singleton et al. 2021) showed that the hospitalized patients correspond to a minority of NHSBT CP donors.

In conclusion, we have for the first time conducted whole-cell and “surfaceome” comparison of the molecular signatures of multiple *ex vivo*-generated macrophages and explored their functional properties. Although these cells are derived from the same founder macrophage population, they display significant and specific alterations in total proteome, surface markers and behavior associated with the specific subtypes generated and their roles in immunity. It should be noted that additional cues such as other cell types or specific niche signals, may also contribute to aspects of polarity which are not captured in our data sets, as a consequence of their generation *ex vivo*. Nevertheless, *t*his work confirms macrophages do not constitute stand-alone phenotypes but instead a continuum of subtypes built on a particular cell theme that are plastic. Importantly, we have dissected M2a and M2c macrophages beyond the reach of previous literature and detail key changes between these subtypes. We also demonstrate that short term exposure to treatment with dexamethasone causes M1 cells to partially repolarize, exhibiting features of both M1 and M2c macrophage subtypes. Additional cues such as other cell types or specific niche signals, may also contribute to aspects of polarity which are not captured in our data sets, as a consequence of their generation *ex vivo*. This work provides evidence and highlights the exciting potential of *ex vivo* culture to recapitulate macrophage polarity, and as a future test bed for screening immune modulation-directed drugs and therapeutics to combat inflammatory disease.

## 5 Conflict of Interest

The authors declare that the research was conducted in the absence of any commercial or financial relationships that could be construed as a potential conflict of interest. The views expressed are those of the authors and not necessarily those of the National Health Service, NIHR, or the Department of Health and Social Care.

## 6 Author Contributions

TCLO, CES and AMT designed the experiments. CES and TCLO conducted the majority of the experimental work. SJC designed and wrote the image analysis software, KJH and MCW performed the proteomics experiments, HEB conducted the serology ELISAs, KR performed the ROS assays, KLHS performed tissue culture experiments and PLM conducted the proteomics analysis and provided insightful discussion. TCLO, CES and AMT wrote the manuscript and all authors read and edited the manuscript prior to submission. CES and AMT contributed equally to conception and supervision of the work. The authors have no conflict of interest to declare.

## 7 Funding

This work was funded by grants from NIHR Blood and Transplant Research Unit in red cell products (IS-BTU-1214-10032), NHS Blood and Transplant (NHSBT) R&D (WP15-05) and a Wellcome Trust PhD Studentship to TCLO (8043 WT 108907/Z/15/Z Dynamic Cell), PLM was supported by a Cancerfonden (Swedish Cancer Society) postdoctoral grant (21 0340 PT) and the work by HEB was supported by the Elizabeth Blackwell Institute (University of Bristol) with funding from the University’s alumni and friends.

## 8 Acknowledgments

The authors would like to thank Dr. Katy Jepson for facilitating the use of the IncuCyte^®^ system and Prof. Imre Berger and Dr. Kapil Gupta for kind gift of SARS-CoV-2 Spike protein. The authors also acknowledge MRC and Wolfson Foundation for establishing the Wolfson Bioimaging Facility (University of Bristol) and for the use of microscopes.

## Notes

https://doi.org/10.5523/bris.11qjhj9l4jpfj2i34kob8d4eug

### Competing Interest Statement

The authors have declared no competing interest.

### Summary of Updates

The authors order has been changed to fully reflect the contributions of the authors. Tiah Oates is now first and Charlotte Severn has moved to second senior author/joint corresponding. Both authors are aware of this and requested this. Contribution section has been adjusted slightly. The front page text and abstract word counts have been corrected Key words have had the numbers removed Font of main text has been changed to arial

