## Supplemental figures for "Characterizing the polarization continuum of macrophage subtypes M1, M2a and M2c"

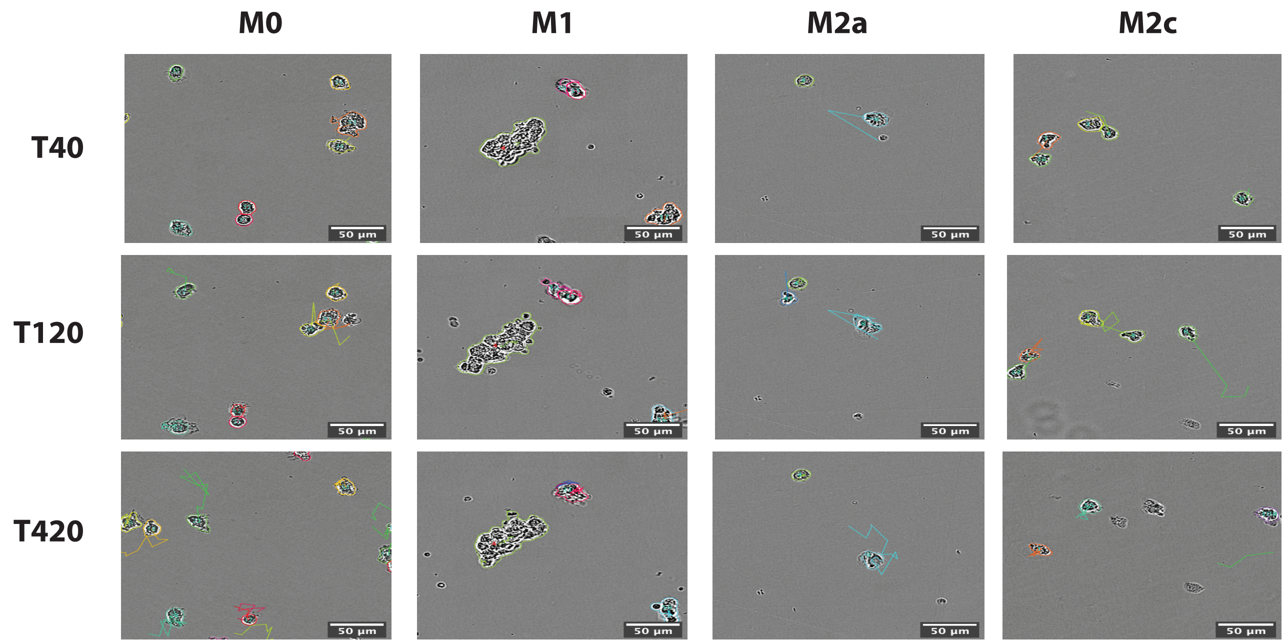


**Supplemental figure 1: Characterisation of cell phenotypes in macrophage polarisation states.** Representative images captured from IncuCyte® analysis at 40 minutes, 120 minutes and 420 minutes. Scale bars are 50 µm.

**A**

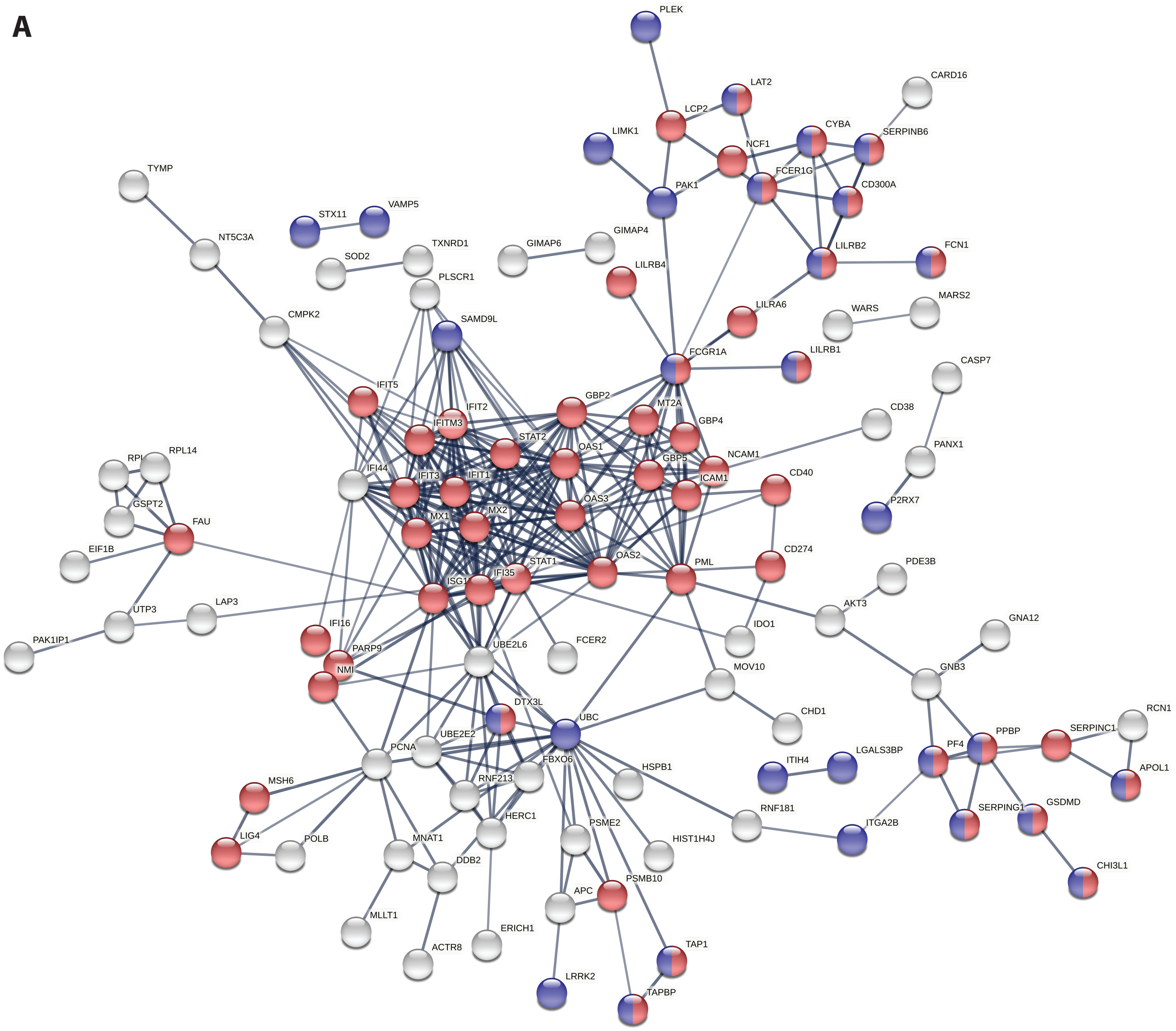


**Supplemental figure 2: Network analysis of whole-cell proteomic data.** A) Network of M1 upregulated proteins where nodes in red denote the gene ontology (GO) term immune response and those in blue vesicle transport.

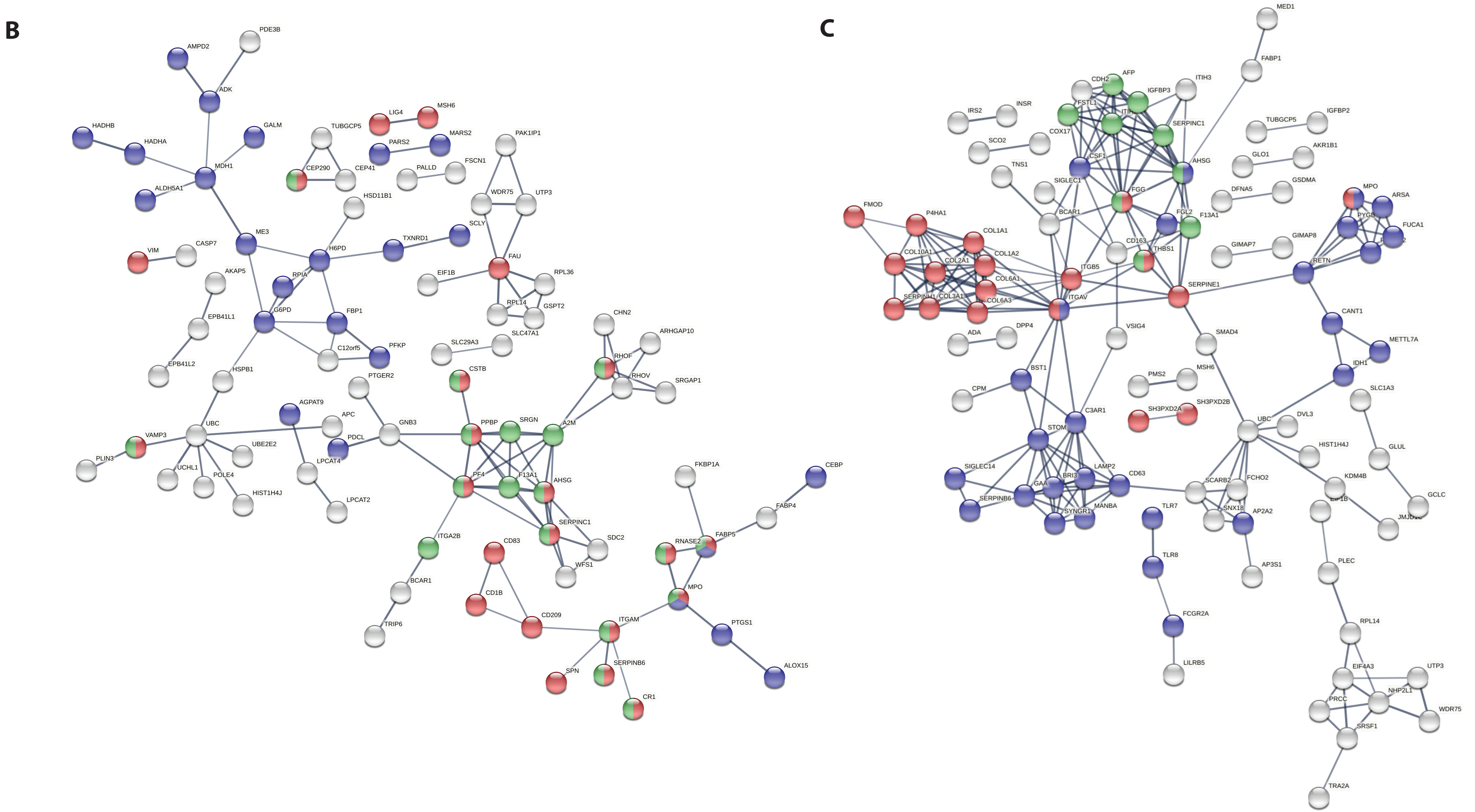


**Supplemental figure 2 continued: Network analysis of whole-cell proteomic data.** B) Upregulated proteins for M2a macrophages where red nodes represent the GO term immune response, green secretion, and blue metabolism. C) M2c upregulated proteins from the following GO terms, where red nodes are extracellular proteins, blue for myeloid leukocyte activation and green for IGF transport and metabolism.

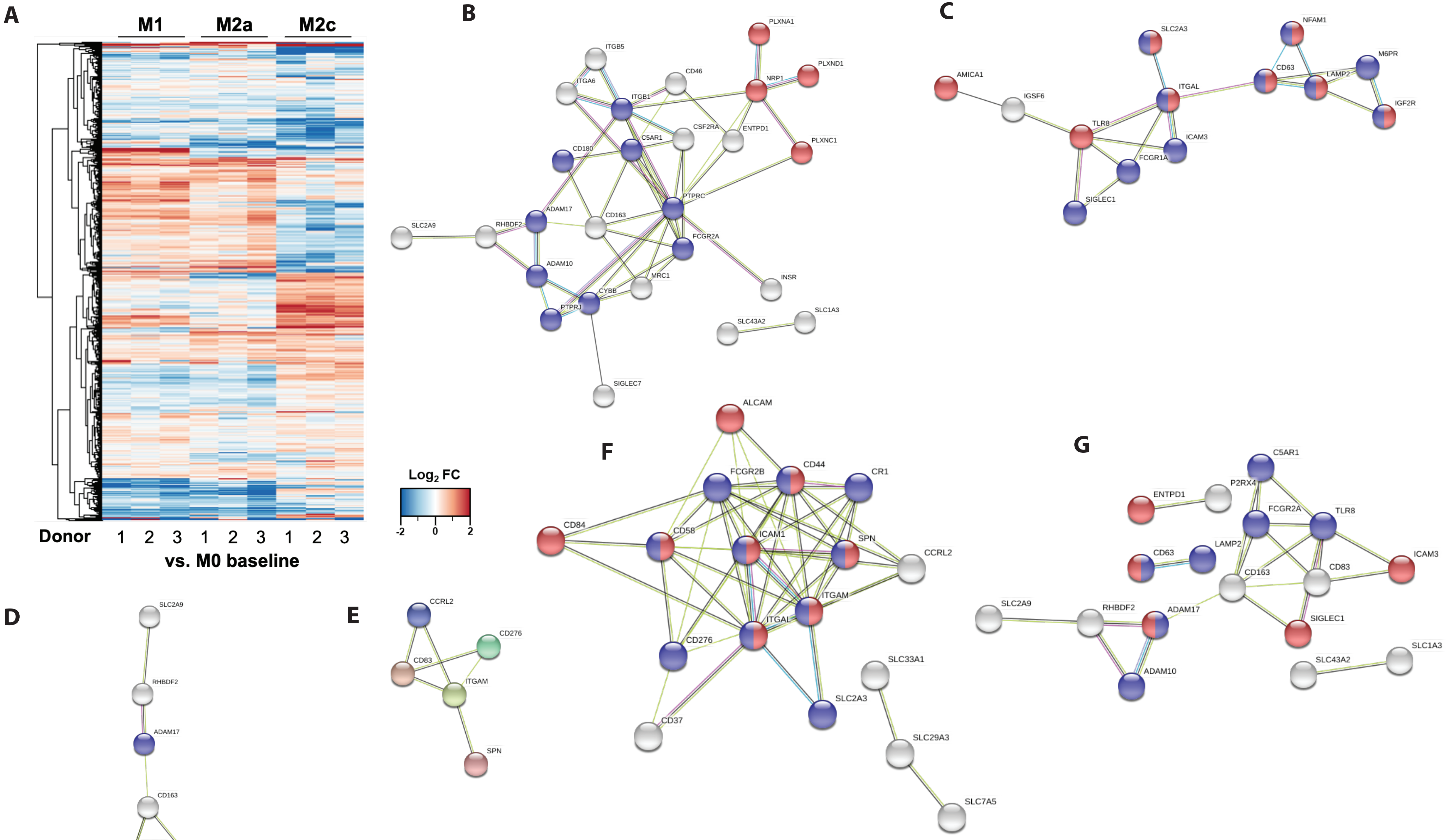


**Supplemental figure 3: Further analysis of surfaceome proteomic dataset.** A) Heatmap visualisation of the TMT surfaceome analysis before filtering for surface proteins only. B-G) STRING visualisation of up and downregulated proteins in macrophage subtypes where node colour responds to the following gene ontology (GO) terms: B) M1 downregulated proteins, red, semaphorin-plexin signalling pathway and blue, leukocyte activation C) M1 upregulated proteins, red, leukocyte activation and blue, vesicle mediated transport D) M2a downregulated proteins, red, semaphoring-plexin signalling pathway and blue, leukocyte mediated immunity E) M2a upregulated proteins F) M2c downregulated proteins, red, cell-cell adhesion, blue, leukocyte activation G) M2c upregulated proteins, red, cell adhesion, blue, leukocyte mediated immunity.

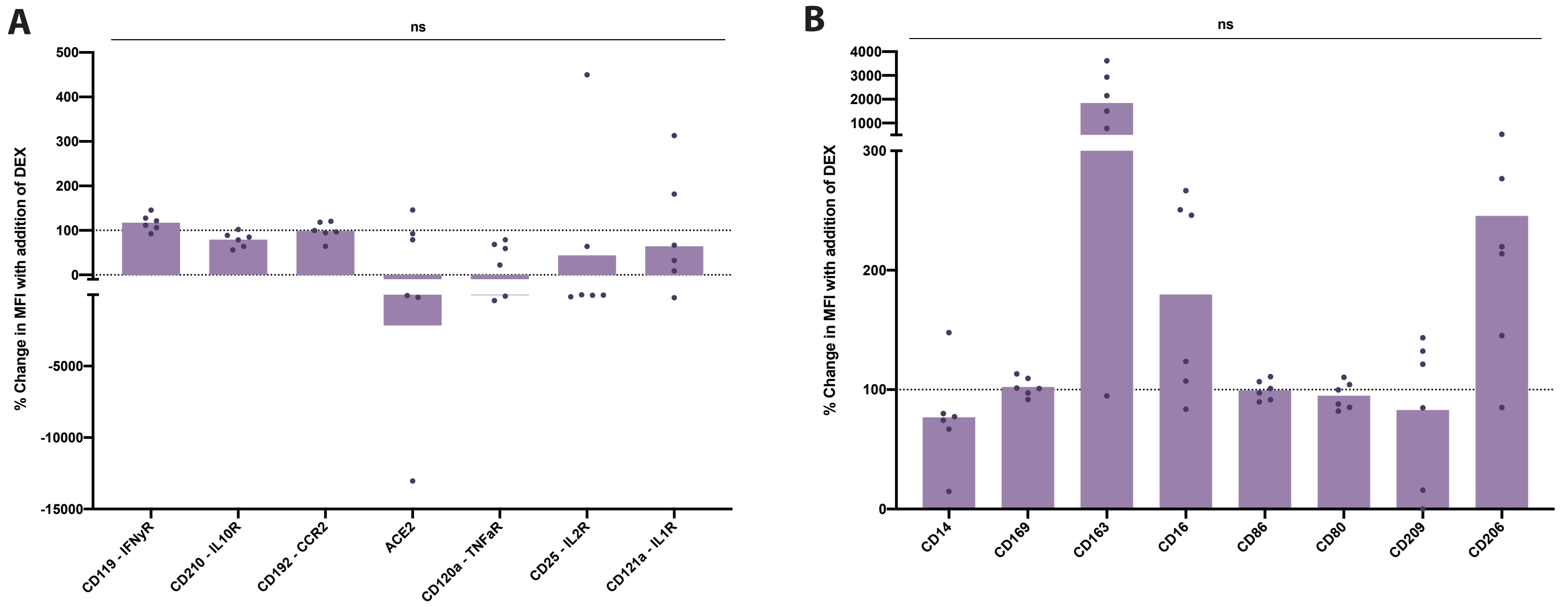


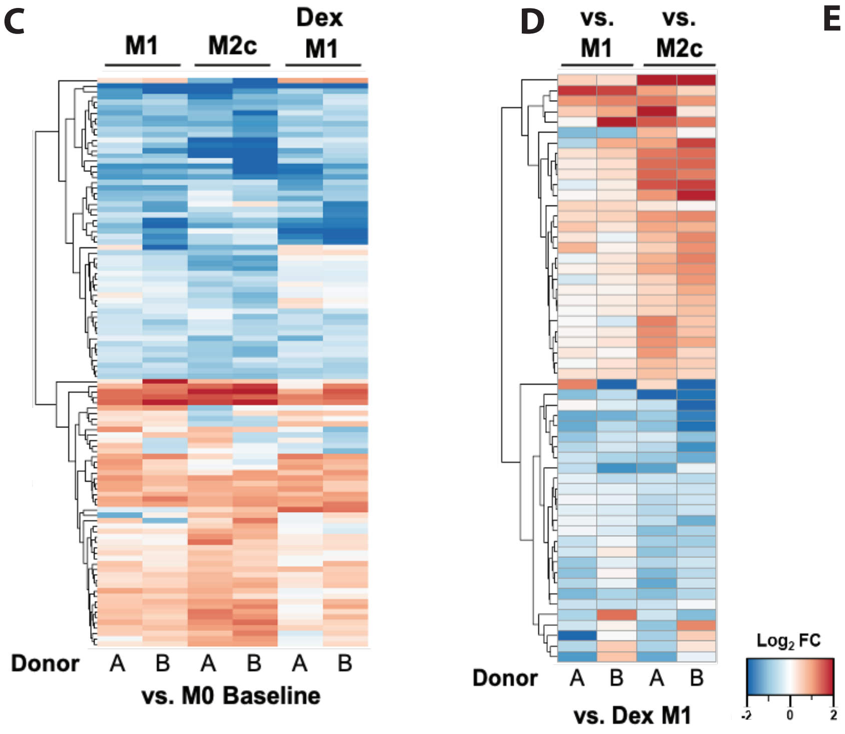


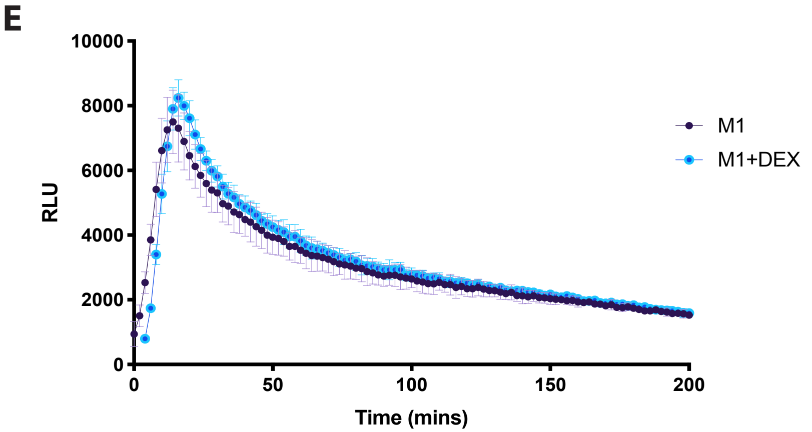


**Supplemental figure 4: Differential expression of surface markers in M1 macrophages after 48 hours treatment with dexamethasone.** A and B) Median fluorescence intensity for flow cytometry presented in Figure 5 a and b. Data presented as a percentage change upon addition of dexamethasone. Dotted line depicts the 100% and subsequently no change in expression. Bar shows mean with each point representing individual samples (N=6, ns $p \geq 0.05$), significance tested using a one sample t-test and Wilcoxon test. C) Heatmap visualisation of TMT surfaceome analysis filtered for surface proteins generated from log2 fold changes in expression in M1, M2c and M1 dexamethasone treated samples with the MØ control as a baseline. Lower expression represented in blue whilst red denotes higher expression. FDR set at 5%. D) Heatmap visualisation of differential expression generated from log2 fold changes in expression of surface markers in M1 and M2c subtypes when using M1 dexamethasone macrophages as a baseline. For C and D) Lower expression represented in blue whilst red denotes higher expression. FDR set at 5%. E) Comparison of respiratory burst formation in M1 macrophages and dexamethasone treated M1 macrophages when stimulated with PMA, N=3.

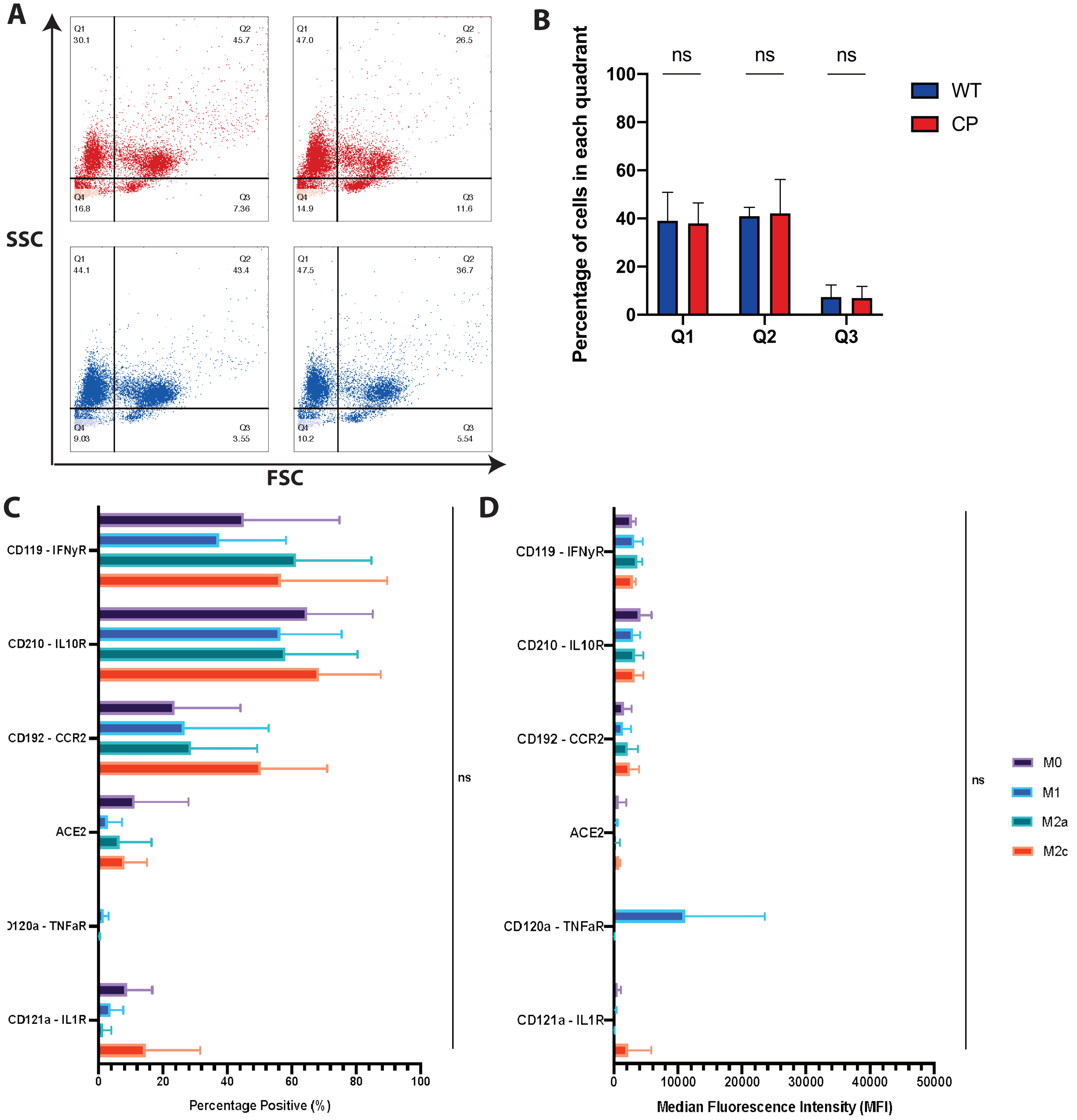


**Supplemental figure 5: Effect of COVID-19 infection on macrophage populations and surface marker expression.** A) Representative FSC/SSC flow cytometry scatter plots demonstrating differences in size and granularity of macrophages isolated from control and CP samples. B) Quantification of percentage of populations for each of the quadrants in A, error bars represent the standard deviation. C) Surface markers in relation to COVID-19 infection tested using flow cytometry on ex vivo generated subtypes. D) Median fluorescence intensity of flow cytometry markers tested in C). All flow cytometry experiments show the mean and standard deviation for the data (N=3,ns $p \geq 0.05$), with a minimum of 10,000 events per sample, significance tested using a one-way ANOVA followed by a Tukeys multiple comparison test, comparing each subtype.
